# Modeling SARS-CoV-2 infection and its individual differences with ACE2-expressing human iPS cells

**DOI:** 10.1101/2021.02.22.432218

**Authors:** Emi Sano, Ayaka Sakamoto, Natsumi Mimura, Ai Hirabayashi, Yukiko Muramoto, Takeshi Noda, Takuya Yamamoto, Kazuo Takayama

## Abstract

Genetic differences are a primary reason for differences in the susceptibility and severity of coronavirus disease 2019 (COVID-19). Because induced pluripotent stem (iPS) cells maintain the genetic information of the donor, they can be used to model individual differences in severe acute respiratory syndrome coronavirus 2 (SARS-CoV-2) infection *in vitro*. Notably, undifferentiated human iPS cells themselves cannot be infected bySARS-CoV-2. Using adenovirus vectors, here we found that human iPS cells expressing the SARS-CoV-2 receptor angiotensin-converting enzyme 2 (ACE2) (ACE2-iPS cells) can be infected with SARS-CoV-2. In infected ACE2-iPS cells, the expression of SARS-CoV-2 nucleocapsid protein, the budding of viral particles, the production of progeny virus, double membrane spherules, and double-membrane vesicles were confirmed. We also evaluated COVID-19 therapeutic drugs in ACE2-iPS cells and confirmed the strong antiviral effects of Remdesivir, EIDD-2801, and interferon-beta. In addition, we performed SARS-CoV-2 infection experiments on ACE2-iPS/ES cells from 8 individuals. Male iPS/ES cells were more capable of producing the virus as compared with female iPS/ES cells. These findings suggest that ACE2-iPS cells can not only reproduce individual differences in SARS-CoV-2 infection *in vitro*, but they are also a useful resource to clarify the causes of individual differences in COVID-19 due to genetic differences.

**Graphical Abstract:** 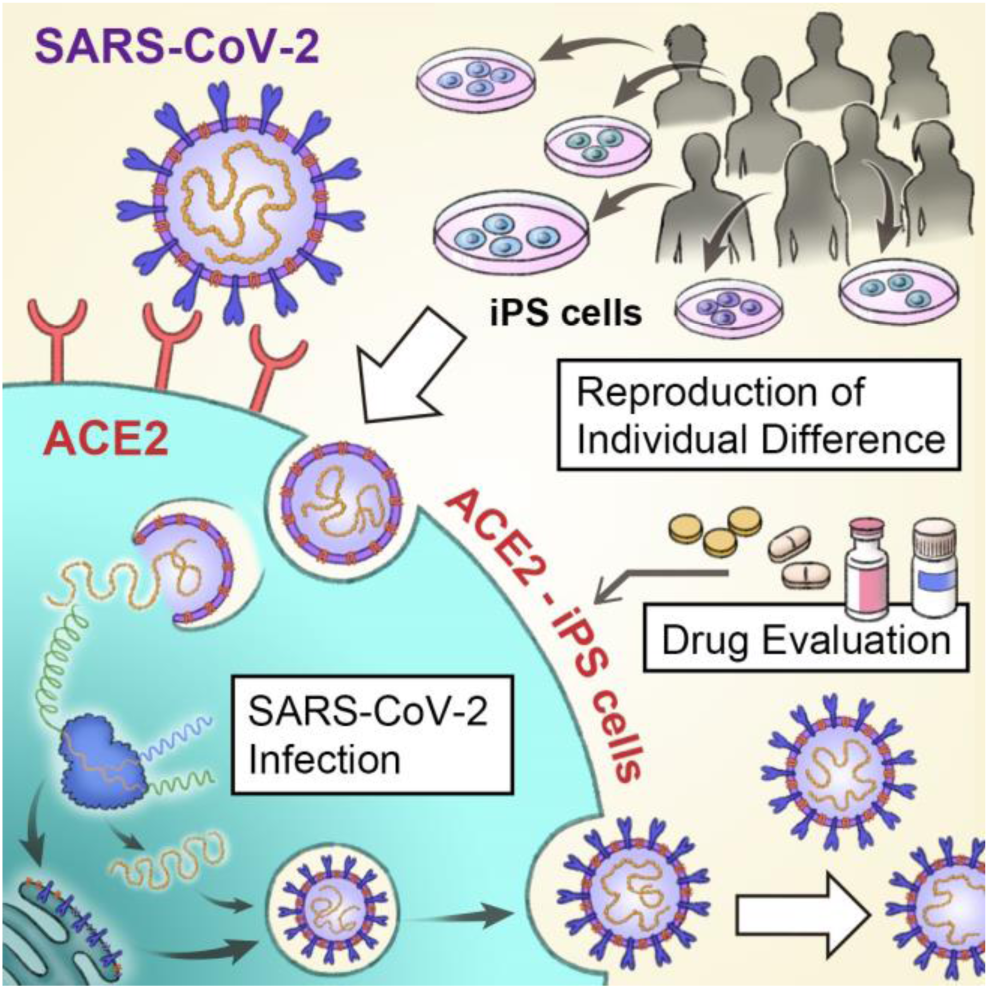

## Introduction

The number of coronavirus disease 2019 (COVID-19) patients and deaths continues to rise. Interestingly, the symptoms of COVID-19 are known to vary widely among individuals and include asymptomatic cases. Genetic differences are one cause for the differences in susceptibility and severity of COVID-19 (Anastassopoulou et al., 2020; Group, 2020; Pairo-Castineira et al., 2020; Zeberg and Pääbo, 2020). Approximately twenty percent of patients infected with severe acute respiratory syndrome coronavirus 2 (SARS-CoV-2) will present severe symptoms (Weiss and Murdoch, 2020). To develop drugs for patients with severe COVID-19, it is necessary to identify the causes of the worsening symptoms. Although several models, including cells such as Vero, Calu-3, and Caco-2 cells, organoids, and animals such as angiotensin-converting enzyme 2 (ACE2)-transgenic mice, hamsters, and ferrets have been used to study SARS-CoV-2 infection (Takayama, 2020), they do not reproduce individual differences well.

Human induced pluripotent stem (iPS) cells can be established from any individual (Takahashi et al., 2007). Furthermore, they are widely used as genetic disease models because they inherit the genetic information of their donors (Park et al., 2008). Our institute (Center for iPS Cell Research and Application, CiRA) has established iPS cells from more than 500 individuals. Because this iPS cell panel was established from a human population of diverse genetic backgrounds, it can be a resource for reproducing individual differences in response to pathogens and drugs. Accordingly, in this study, we attempted to reproduce the SARS-CoV-2 infection and its individual differences using this panel.

It has been reported that SARS-CoV-2 infection is dependent on the expression of ACE2 and transmembrane protease, serine 2 (TMPRSS2) in host cells (Hoffmann et al., 2020). Type II alveolar epithelial cells, bronchial ciliated cells, pharyngeal epithelial cells, and intestinal epithelial cells all express high levels of ACE2 (Ziegler et al., 2020) and can be easily infected by SARS-CoV-2. It has also been reported that embryonic stem (ES) cell-derived type II alveolar epithelial cells and intestinal epithelial cells can be infected with SARS-CoV-2 (Han et al., 2021). However, the investigation of infection differences at the individual level and large scale using iPS cell-derived somatic cells is hindered by the long time (generally more than 3 weeks) to differentiate the iPS cells and the variable differentiation efficiency among iPS cell lines (Kajiwara et al., 2012). SARS-CoV-2 infection experiments in undifferentiated iPS cells would reduce the time.

However, due to the low expression of ACE2 and TMPRSS2, SARS-CoV-2 does not infect iPS cells. Therefore, in this study, we identified SARS-CoV-2-related genes that enable SARS-CoV-2 to infect undifferentiated human iPS cells. To facilitate gene overexpression experiments with multiple iPS cell lines, we used adenovirus (Ad) vectors. Because the gene transfer efficiency of Ad vectors to undifferentiated iPS cells is almost 100%, there was no need to obtain clones expressing SARS-CoV-2-related genes. Next, we investigated whether the life cycle of SARS-CoV-2 can be reproduced in iPS cells expressing these SARS-CoV-2-related genes. We also show the value of ES/iPS cells to test drugs and gender effects. Our findings indicate that iPS cells expressing SARS-CoV-2-related genes can be a useful to study SARS-CoV-2 infection and its individual patient differences.

## Results

### SARS-CoV-2 does not infect undifferentiated iPS cells

First, we examined whether undifferentiated iPS cells could be infected by SARS-CoV-2 (**Fig. S1A**). Before conducting this experiment, we examined the expression levels of viral receptors and proteases in undifferentiated iPS cells (**Fig. S1B**). The gene expression level of *ACE2* was low, but that of *CD147* was high. CD147 is reported as a coronavirus receptor (Wang et al., 2020). TMPRSS2, a protease, was also expressed in undifferentiated iPS cells. We thus tried to infect undifferentiated iPS cells with SARS-CoV-2, but the morphology of the iPS cell colonies did not change (**Fig. S1C**). In addition, virus genome in the cell culture supernatant (**Fig. S1D**) and the production of infectious virus (**Fig. S1E**) were not detected. The gene expression levels of undifferentiated markers (**Fig. S2A**) and innate immune response-related markers (**Fig. S2B**) were also unchanged. Furthermore, the expression of SARS-CoV-2 nucleocapsid (N) protein was not detected (**Fig. S2C**). Together, these results indicated that SARS-CoV-2 does not infect undifferentiated iPS cells.

### ACE2 expression is required for SARS-CoV-2 to infect human iPS cells

Because human ACE2 and TMPRSS2 are known to be important for SARS-CoV-2 to infect cells, we overexpressed human ACE2 and TMPRSS2 in undifferentiated iPS cells by using Ad vectors (**Fig. 1A**). The overexpression of ACE2 in iPS cells (ACE2-iPS cells) caused a large amount of SARS-CoV-2 infection (**Fig. 1B**). Additionally, the amount of virus genome in the cell culture supernatant increased (**Fig. 1C**). This was not the case if only overexpressing TMPRSS2. Furthermore, two days after the ACE2-iPS cells were infected with SARS-CoV-2, cell fusion was observed (**Fig. 1D**), and four days after many of the cells died. Therefore, these results indicate that ACE2 expression is required for SARS-CoV-2 to infect undifferentiated iPS cells.

**Figure 1.**
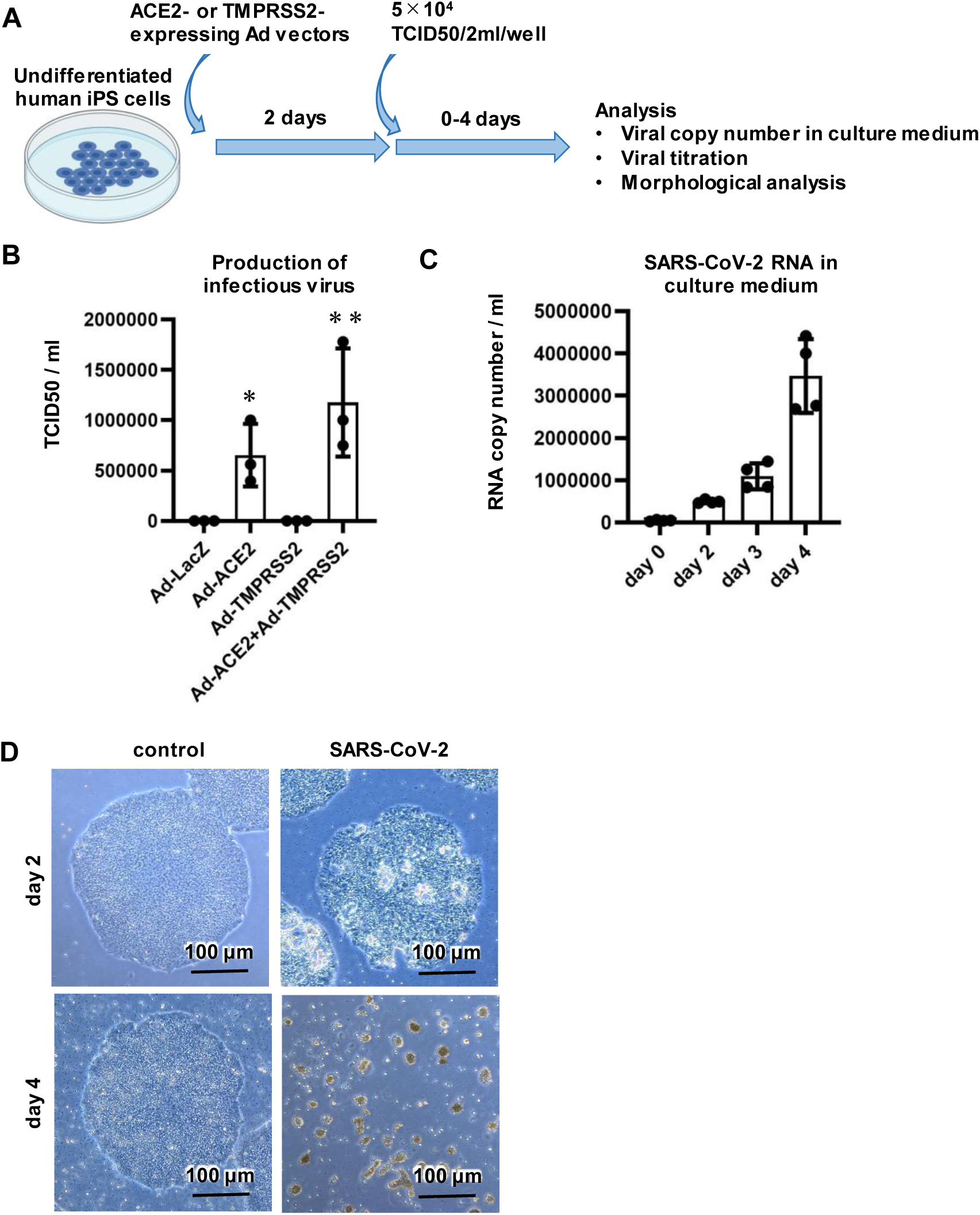
Efficient SARS-CoV-2 infection and replication in ACE2-iPS cells. (**A**) Undifferentiated human iPS cells (1383D6) were transduced with 600 vector particles (VP)/cell of LacZ-, ACE2-, or TMPRSS2-expressing Ad vectors (Ad-LacZ, Ad-ACE2, or Ad-TMPRSS2, respectively) for 2 hr and then cultured with AK02 medium for 2 days. ACE2-expressing human iPS (ACE2-iPS) cells were infected with SARS-CoV-2 (5×10^4^ TCID50/well) for 2 hr and then cultured with AK02 medium. (**B**) The amount of infectious virus in the supernatant was measured by the TCID50 assay. One-way ANOVA followed by Tukey’s post hoc test (\**p*<0.05, \*\**p*<0.01, compared with Ad-LacZ). (**C**) At days 0, 2, 3, and 4 after the SARS-CoV-2 infection, the viral RNA copy number in the cell culture supernatant was measured by qPCR. (**D**) At days 2 and 4 after the SARS-CoV-2 infection, phase images of infected ACE2-iPS cells were obtained. Data are shown as means ± SD (*n*=3).

### Characterization of SARS-CoV-2-infected ACE2-iPS cells

Transmission electron microscope (TEM) images of ACE2-iPS cells infected with SARS-CoV-2 were obtained (**Figs. 2 and S3**). Zippered endoplasmic reticulum (**Fig. 2B**), double-membrane spherules (DMS) (**Fig. 2B**) (Maier et al., 2013), and viral particles near the cell membrane (black arrow) (**Fig. 2D**) were observed. So was an endoplasmic reticulum-Golgi intermediate compartment (ERGIC) containing SARS-CoV-2 particles (black arrows) (**Figs. 2C, S3**). Double membrane vesicles (DMVs, black arrows) were also observed in infected ACE2-iPS cells (**Fig. S3**). DMVs are the central hubs for viral RNA synthesis (Klein et al., 2020). These structures were not observed in the uninfected ACE2-iPS cells. These TEM images show that the life cycle of SARS-CoV-2 can be observed in ACE2-iPS cells.

**Figure 2.**
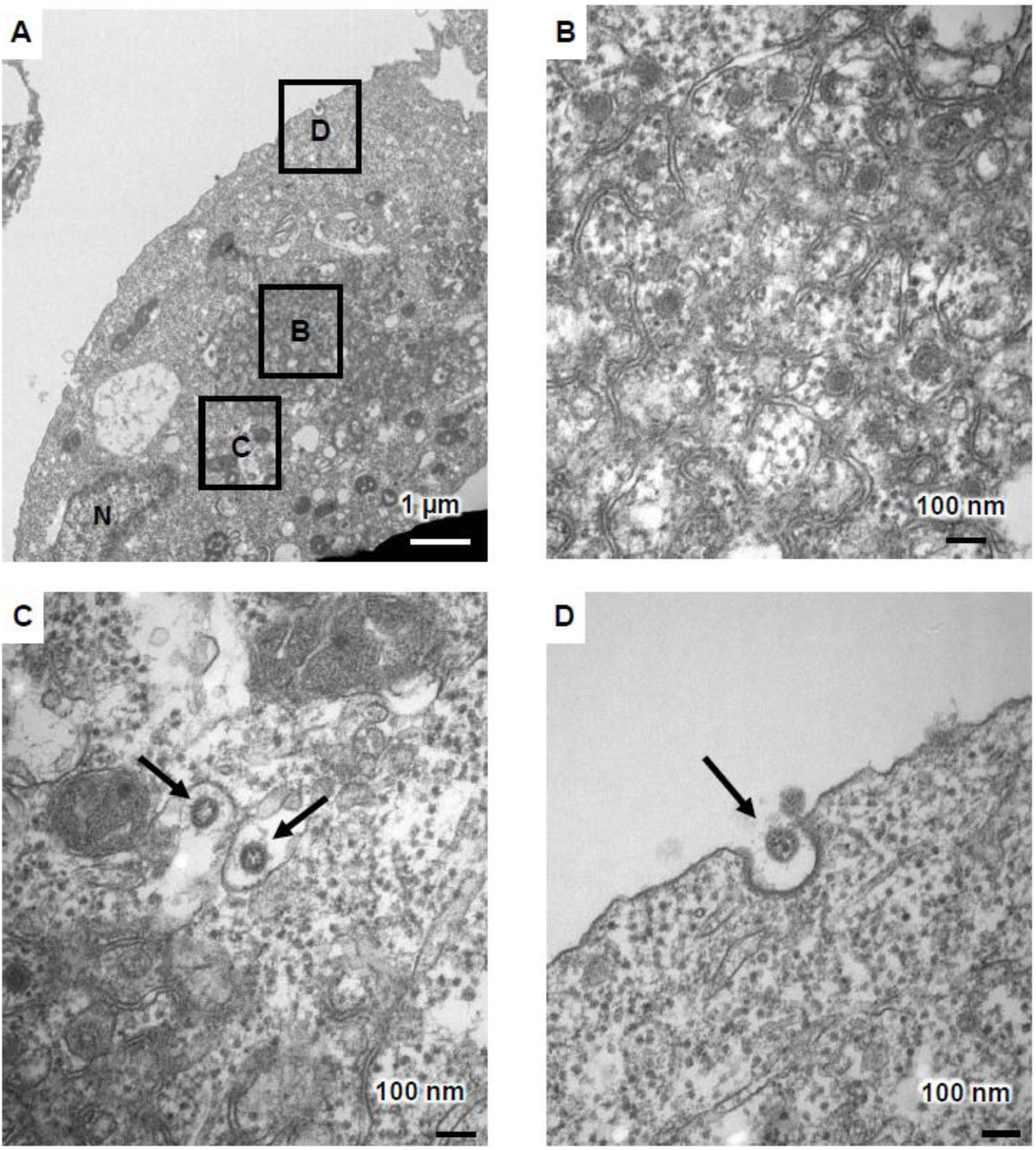
TEM images of infected ACE2-iPS cells. (**A**) TEM images of infected ACE2-iPS cells. Zippered endoplasmic reticulum, double-membrane spherule (DMS) (**B**), and virus particles in the ERGIC (black arrows) (C) and near the cell membrane (black arrows) (**D**) were observed.

Next, we analyzed gene and protein expressions in uninfected and infected iPS cells 3 days after the viral infection. The intracellular viral genome and *ACE2* expression levels in ACE2-iPS cells infected with SARS-CoV-2 were high (**Fig. 3A**). At the same time, ACE2 overexpression and SARS-CoV-2 infection did not alter the gene expression levels of undifferentiated markers (**Fig. 3B**) or innate immune response-related markers (**Fig. 3C**). The gene expression levels of endoderm markers except for *CER1* (**Fig. S4A**) and SARS-CoV-2-related genes (*CD147*, *NRP1*, and *TMPRSS2*) (**Figs. S4B, 3A**) were also unchanged. Immunostaining data showed that SARS-CoV-2 N protein was strongly expressed in ACE2-iPS cells 2 days after the infection (**Figs. 3D, S5**).

**Figure 3.**
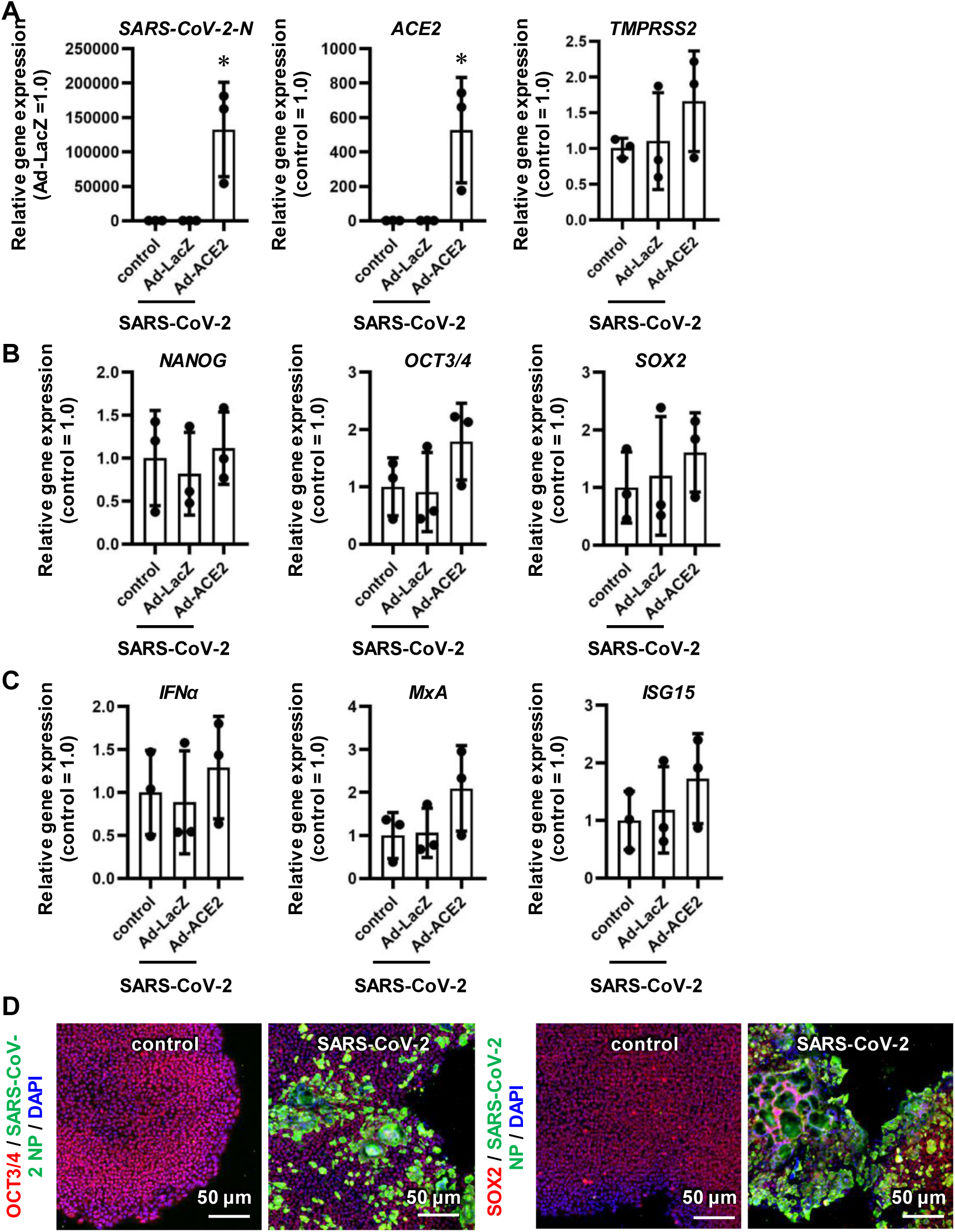
The pluripotent state of ACE2-iPS cells is not affected by SARS-CoV-2 infection. LacZ- or ACE2-expressing human iPS cells (LacZ-iPS cells and ACE2-iPS cells, respectively) were infected with SARS-CoV-2 (5×10^4^ TCID50/well) for 2 hr and then cultured with AK02 medium for 2 or 3 days. Control human iPS cells were not transduced with Ad vectors. (**A**) The gene expression levels of viral genome, *ACE2*, and *TMPRSS2* were examined by qPCR analysis. (**B**) The gene expression levels of pluripotent markers (*NANOG*, *OCT3/4*, and *SOX2*) and (**C**) innate immunity-related markers (*IFNα*, *MxA*, and *ISG15*) were examined by qPCR. (**D**) Immunofluorescence analysis of SARS-CoV-2 NP (green), OCT3/4 (red), and SOX2 (red) in uninfected and infected ACE2-iPS cells. Nuclei were counterstained with DAPI (blue). One-way ANOVA followed by Tukey’s post hoc test (\**p*<0.05, compared with Ad-LacZ). Data are shown as means ± SD (*n*=3).

We also performed RNA-seq analysis in uninfected and infected ACE2-iPS cells. The colored dots in the volcano plot in **figure 4A** indicate genes whose expression levels changed significantly more than 4-fold. In total, this change occurred in 6.7% of all genes (**Fig. S6A**). A GO term analysis was performed for these genes (**Figs. S6B, S6C**). None of the genes included undifferentiated markers (**Fig. 4B**) or innate immune-response markers (**Fig. 4C**). The gene expression levels of ectoderm, mesoderm, and endoderm markers were also unchanged after infection with SARS-CoV-2 (**Fig. S7**). Overall, these results suggest that human iPS cells maintain an undifferentiated state even when SARS-CoV-2 replicates in large numbers.

**Figure 4.**
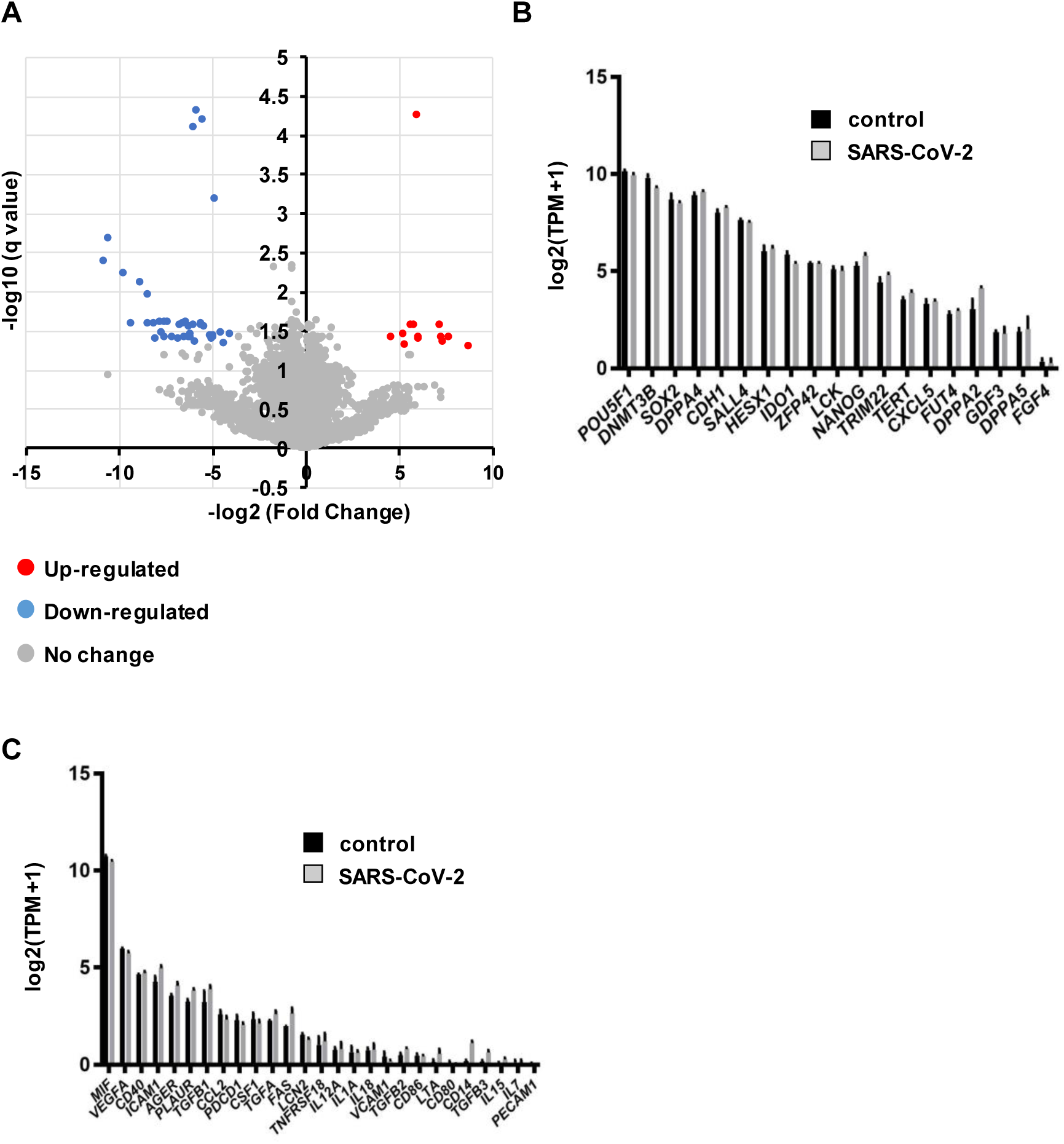
Global gene expression analysis in infected ACE2-iPS cells. RNA-seq analysis was performed in uninfected or infected ACE2-iPS cells. (**A**) A volcano plot of uninfected ACE2-iPS cells vs. infected ACE2-iPS cells. (**B, C**) Bar plots of pluripotent genes (**B**) and immune-related genes (**C**) in uninfected ACE2-iPS cells (control) and infected ACE2-iPS cells (SARS-CoV-2) are shown.

### Evaluation of COVID-19 candidate drugs using ACE2-iPS cells

Next, we examined whether ACE2-iPS cells could be used for drug screening. We tried eight drugs used in COVID-19 clinical trials. Vero cells were used as the control. After exposing the cells to various concentrations of a drug, the number of viral RNA copies in the culture supernatant was quantified (**Fig. 5A**). Data fitting resulted in sigmoid curves, and half-maximal effective concentrations (EC50) were calculated (**Fig. 5B**). Among the eight drugs, the antiviral effect of Remdesivir was strongest. On the other hand, Chloroquine and Favipiravir did not inhibit viral replication, and Ivermectin was highly cytotoxic (**Fig. S8A**). The EC50 and half-maximal cytotoxic concentration (CC50) values of Ivermectin were almost the same between control and infected ACE2-iPS cells (**Fig. S8B**). With the exception of interferon-beta, drug effects were stronger in ACE2-iPS cells than in Vero cells. Lastly, we confirmed the anti-viral effects of RNA-dependent RNA Polymerase (RdRp) inhibitors (Remdesivir and EIDD-2801) and TMPRSS2 inhibitors (Camostat and Nafamostat) in ACE2-iPS cells, indicating that ACE2-iPS cells can be used to evaluate COVID-19 drug candidates.

**Figure 5.**
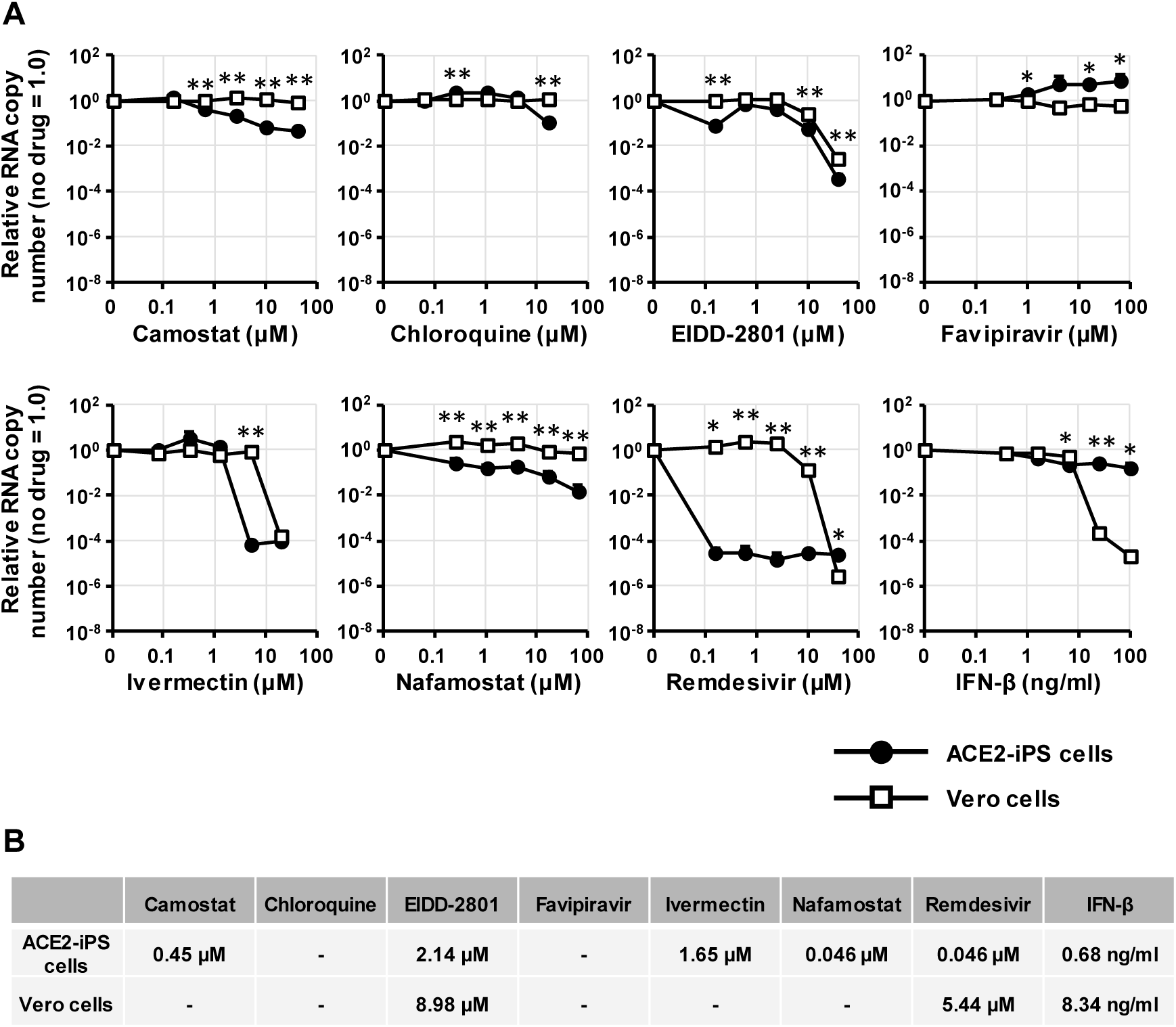
Evaluation of anti-COVID-19 drugs in ACE2-iPS cells. (**A**) ACE2-iPS cells and Vero cells (2.0×10^4^ cells/well) were infected with SARS-CoV-2 (2.0×10^3^ TCID50/well) in the presence or absence of a drug and then cultured with medium for 4 and 2 days, respectively. The viral RNA copy number in the cell culture supernatant was measured by qPCR. (**B**) The EC50 values (μM) of anti-COVID-19 drugs in ACE2-iPS cells and Vero cells were calculated using GraphPad Prism8 and are summarized. Data are shown as means ± SD (*n*=3). Unpaired two-tailed Student’s *t*-test (\**p*<0.05, \*\**p*<0.01; ACE2-iPS vs Vero cells).

### SARS-CoV-2 infection experiments in various ACE2-iPS/ES cell lines

Finally, we performed SARS-CoV-2 infection experiments using human ACE2-iPS/ES cells established from eight donors. The replication efficiency of the virus was different among the ACE2-iPS/ES cell lines (**Fig. 6A**). Note that there was no significant difference in *ACE2* expression levels in the ACE2-iPS/ES cell lines (**Fig. S9**). Interestingly, the viral replication capacity of male ACE2-iPS/ES cells was higher than that of female ACE2-iPS/ES cells (**Fig. 6B**), suggesting that sex differences in the susceptibility to SARS-CoV-2 can be reproduced using ACE2-iPS/ES cells. Recently, it has been speculated that the expression levels of *androgen receptor* and its target gene, *TMPRSS2*, are involved in the sex differences in the SARS-CoV-2 infection (Wambier et al., 2020). The *TMPRSS2* expression levels appeared to be higher in male iPS/ES cells than in female iPS/ES cells (**Fig. 6C**), but there was no significant difference (**Fig. 6D**).

**Figure 6.**
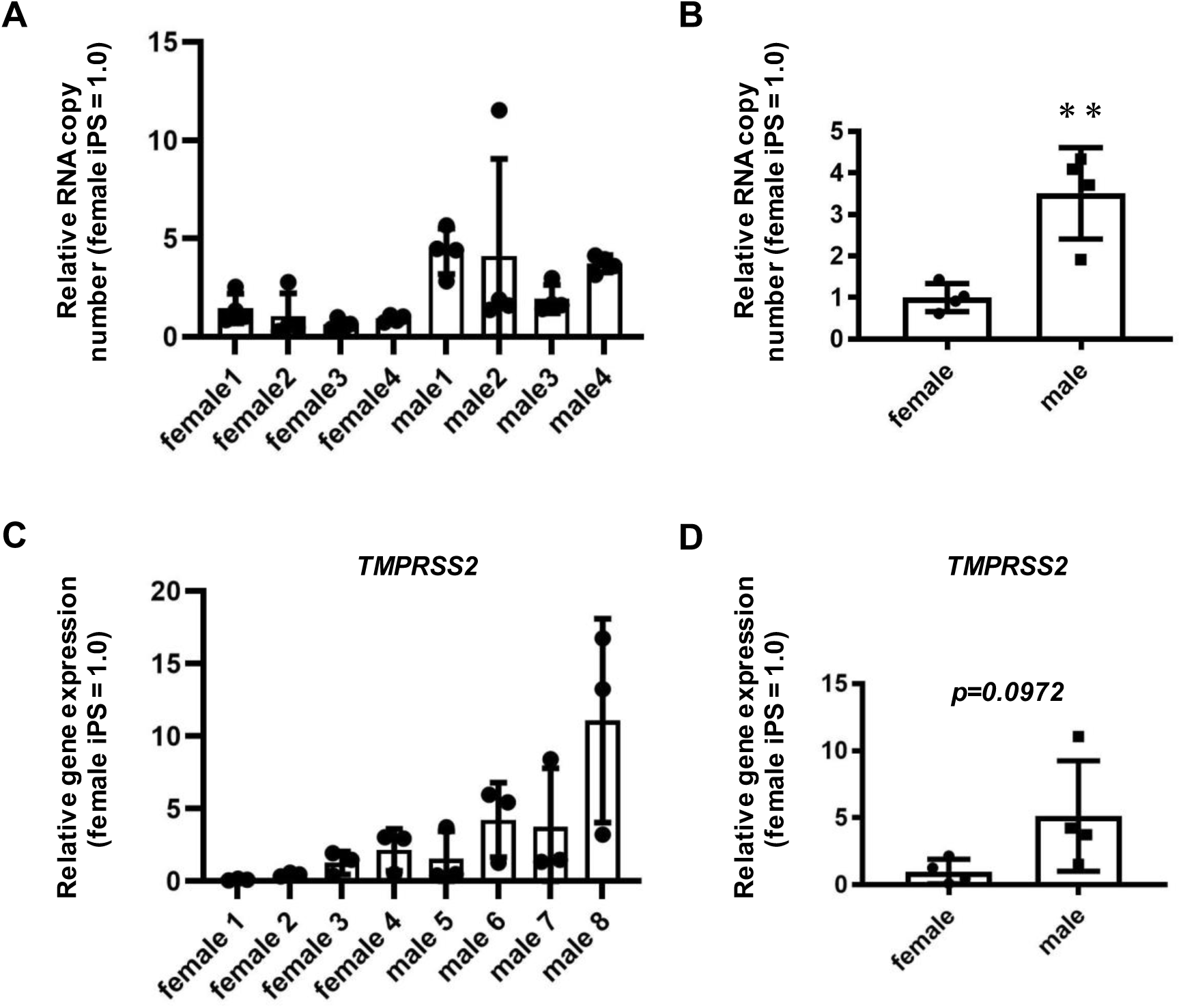
Sex differences of the SARS-CoV-2 infection rate in ACE2-ES/iPS cells. Four female ES/iPS cell lines and four male ES/iPS cell lines were transduced with 600 VP/cell of ACE2-expressing Ad vectors (Ad-ACE2) for 2 hr and then cultured with AK02 medium for 2 days. The cells were then infected with SARS-CoV-2 (5×10^4^ TCID50/well) for 2 hr and cultured with AK02 medium. (**A**) The viral RNA copy number in the cell culture supernatant was measured by qPCR for each cell line. (**B**) The viral RNA copy number in the cell culture supernatant was compared between female iPS/ES cells and male iPS/ES cells. (**C**) The *TMPRSS2* expression levels were measured by qPCR for each cell line. (**D**) The *TMPRSS2* expression levels were compared between female iPS/ES cells and male iPS/ES cells. Unpaired two-tailed Student’s *t*-test (\*\**p*<0.01). Data are shown as means ± SD (*n*=3). Female1:H9, Female2: KhES1, Female3:KhES2, Female4: 201B7, Male1:H1, Male2:KhES3, Male3:Tic, Male4:1383D6.

## Discussion

In this study, we showed that the life cycle of SARS-CoV-2 can be reproduced in human iPS cells overexpressing ACE2. In addition, we were able to confirm the effects of two TMPRSS2 inhibitors (Camostat and Nafamostat) and two RdRp inhibitors (Remdesivir and EIDD-2801) using these ACE2-iPS cells. Finally, we showed a difference in the efficiency of infection of SARS-CoV-2 among ACE2-iPS/ES cells from 8 donors. These results suggest that by using our iPS cell panel, it will be possible to investigate the effects of race and blood type as well as gender on SARS-CoV-2 infection. By conducting SARS-CoV-2 infection experiments using a panel of iPS cells for which genomic information has been obtained, it will also be possible to find genomic mutations that appear with high frequency in susceptible cells.

We observed a difference in the infection efficiency of ACE2-iPS/ES cells between donors, but we do not know if this difference reflects the sensitivity of the original donors to COVID-19, since none were COVID-19 patients. Therefore, we are currently establishing iPS cells from severe and mild COVID-19 patients. We plan to conduct infection experiments in the newly established iPS cells and compare the results with the symptoms of the original patients. These experiments could clarify whether the results of SARS-CoV-2 infection experiments in iPS cells reflect the symptoms of the original donors.

Recently, GWAS analyses was performed on severe and mild COVID-19 patients (Group, 2020; Pairo-Castineira et al., 2020). The analysis of the severe COVID-19 GWAS group indicated that mutations in the SLC6A20, LZTFL1, CCR9, FYCO1, CXCR6, and XCR1 genes are associated with the severity of the symptoms (Group, 2020). Pairo-Castineira et al. suggested that the mutations and expression levels of the IFNAR2 and TYK2 genes are also associated with the severity of COVID-19 (Pairo-Castineira et al., 2020). However, the function of these genes in COVID-19 is still not fully understood. Further analysis can be performed by examining the relationship between viral infection and gene mutations and expressions in our iPS cell panel. Because single nucleotide mutations can be easily introduced into iPS cells (Kim et al., 2018), the function of these mutations can be studied using genome-edited iPS cells.

To infect iPS cells with SARS-CoV-2, we overexpressed ACE2. However, if ACE2 and its related genes are responsible for the individual differences in SARS-CoV-2 infection, our system will not be effective. To analyze the mutation and expression of ACE2 and its related genes, it is essential to use somatic cells expressing ACE2. Also, our system is not effective for non-genetic causes of individual differences in COVID-19 severity. For example, it has been speculated that age-related differences in COVID-19 symptoms are due to impaired cytotoxic CD8+T cell responses (Westmeier et al., 2020). For such studies, it is more suitable to use blood samples.

Accordingly, there may be multiple causes of the individual differences in COVID-19 symptoms. The ACE2-iPS cells that we have developed in this study will be one tool to elucidate the cause of these individual differences, which will help identify vulnerable populations and develop new drugs.

## Abbreviations

Ad: adenovirus
ACE2: angiotensin-converting enzyme 2
COVID-19: coronavirus disease 2019
DMS: double membrane spherule
DMV: double membrane vesicle
ES cells: embryonic stem cells
ERGIC: endoplasmic reticulum-Golgi intermediate compartment
GWAS: genome-wide association studies
CC50: half maximal cytotoxic concentration
EC50: half maximal effective concentration
iPS cells: induced pluripotent stem cells
N: nucleocapsid
PBMCs: peripheral blood mononuclear cells
RdRp: RNA-dependent RNA Polymerase
SARS-CoV-2: severe acute respiratory syndrome coronavirus 2
TMPRSS2: transmembrane protease, serine 2
TEM: transmission electron microscope
VP: vector particles

## Acknowledgements

The SARS-CoV-2 strain used in this study (SARS-CoV-2/Hu/DP/Kng/19-027) was kindly provided by Dr. Tomohiko Takasaki and Dr. Jun-Ichi Sakuragi (Kanagawa Prefectural Institute of Public Health). **Figures 1A, S1A** were created using Biorender (https://biorender.com). We thank Dr. Misaki Ouchida (Kyoto University) for creating the graphical abstract, Dr. Peter Karagiannis (Kyoto University) for critical reading of the manuscript, Dr. Masato Nagagawa (Kyoto University) for providing the human iPS cells, Dr. Yoshio Koyanagi and Dr. Kazuya Shimura (Kyoto University) for the setup and operation of the BSL-3 laboratory at Kyoto University, Dr. Toru Okamoto (Osaka University), Dr. Akatsuki Saito (University of Miyazaki), Dr. Hirofumi Ohashi, and Dr. Koichi Watashi (National Institute of Infectious Diseases) for helpful discussion, and Ms. Kazusa Okita and Ms. Satoko Sakurai (Kyoto University) for technical assistance with the RNA-seq experiments. This research was supported by the iPS Cell Research Fund, the COVID-19 Private Fund (to the Shinya Yamanaka laboratory, CiRA, Kyoto University), and the Japan Agency for Medical Research and Development (AMED) (20fk0108263s0201).

## Author Contributions

ES performed the SARS-CoV-2 experiments, analyses, and statistical analysis

AS performed the human cell culture and qPCR analyses

NM performed the drug experiments

AH obtained the TEM images

YM obtained the TEM images

TN obtained the TEM images

TY performed the RNA-seq analysis

KT performed the SARS-CoV-2 experiments and analyses and wrote the paper

---

**Figure S1.**
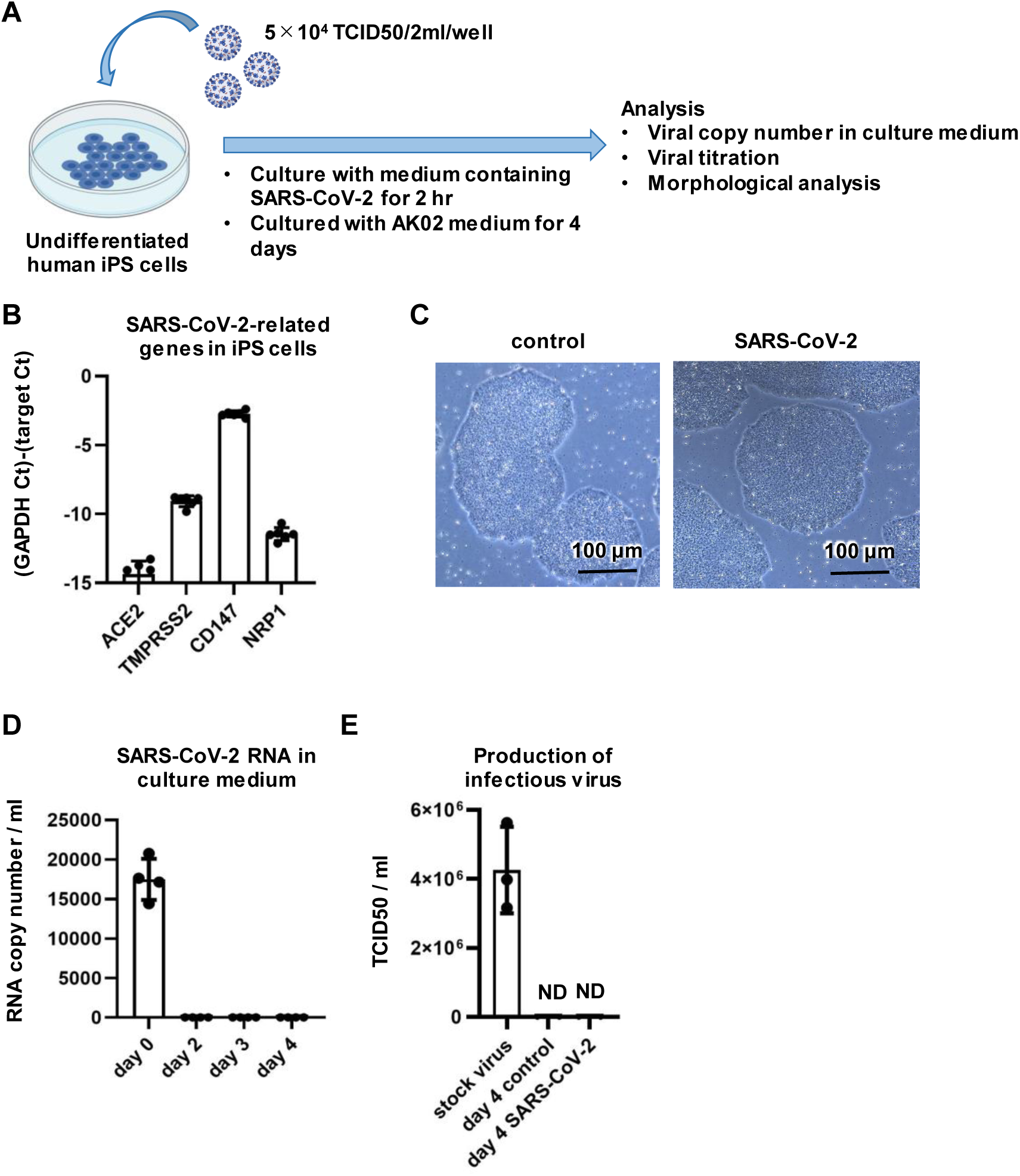
SARS-CoV-2 cannot infect human iPS cells. (**A**) Undifferentiated human iPS cells (1383D6) were infected with SARS-CoV-2 (5×10^4^ TCID50/well) for 2 hr and then cultured with AK02 medium for 4 days. (**B**) The gene expression levels of *ACE2*, *TMPRSS2*, *CD147*, and *NRP1* in human iPS cells were examined by qPCR. (**C**) Phase images of uninfected and infected human iPS cells are shown. (**D**) At days 0, 2, 3 and 4 after the infection, the viral RNA copy number in the cell culture supernatant was measured by qPCR. day 0 = medium containing initial virus (5×104 TCID50/2ml). (**E**) The amount of infectious virus in the supernatant was measured by the TCID50 assay. Data are shown as means ± SD (*n*=3). ND=not detected.

**Figure S2.**
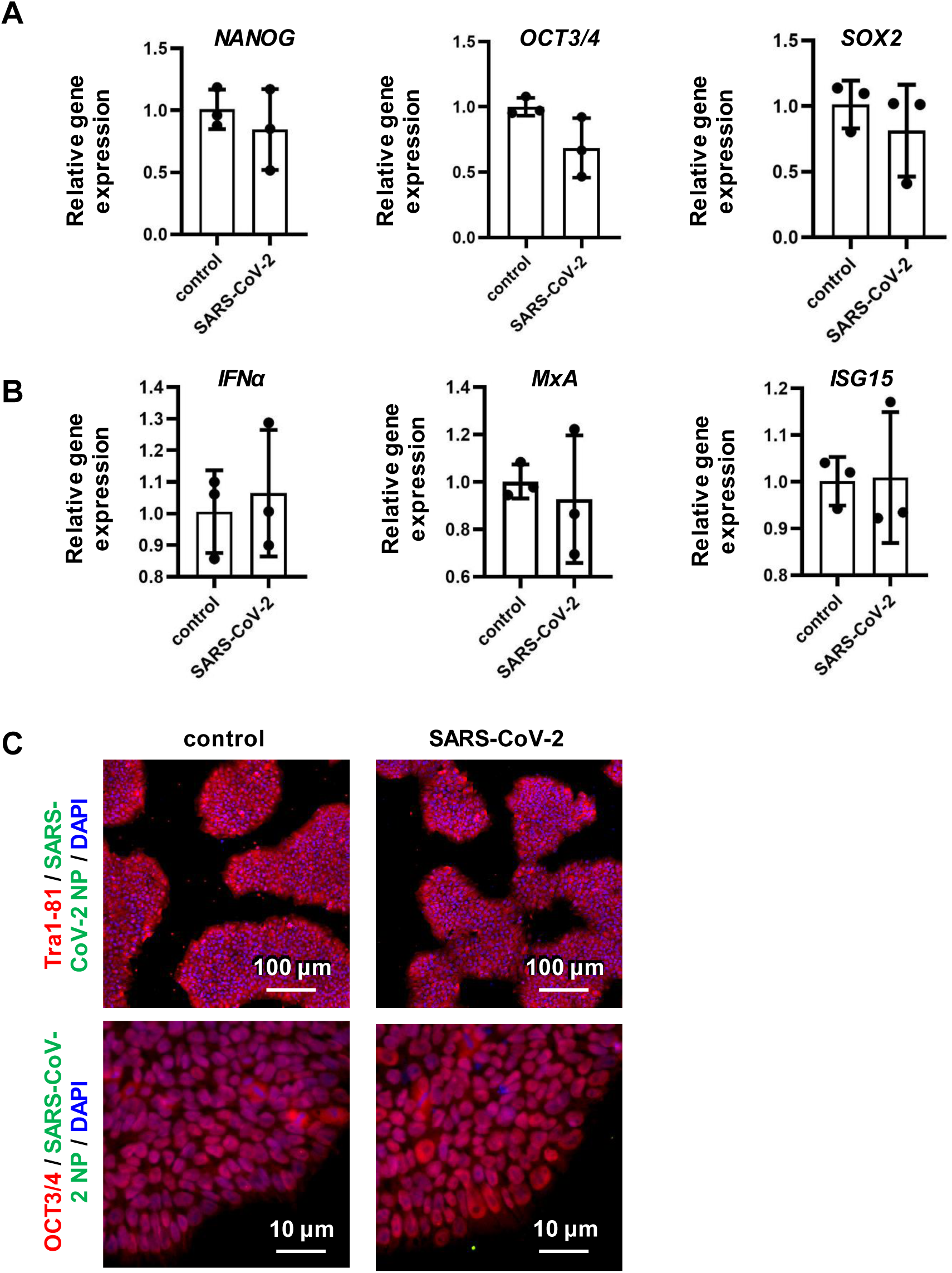
The pluripotent state of human iPS cells is not affected by SARS-CoV-2 infection. The gene expression levels of pluripotent markers (*NANOG*, *OCT3/4*, and *SOX2*) (**A**) and innate immunity-related markers (*IFNα*, *MxA*, and *ISG15*) (**B**) in uninfected and infected human iPS cells were examined by qPCR. (**C**) Immunofluorescence analysis of SARS-CoV-2 NP (green), Tra1-81 (red), and OCT3/4 (red) in uninfected and infected human iPS cells. Nuclei were counterstained with DAPI (blue). Data are shown as means ± SD (*n*=3).

**Figure S3.**
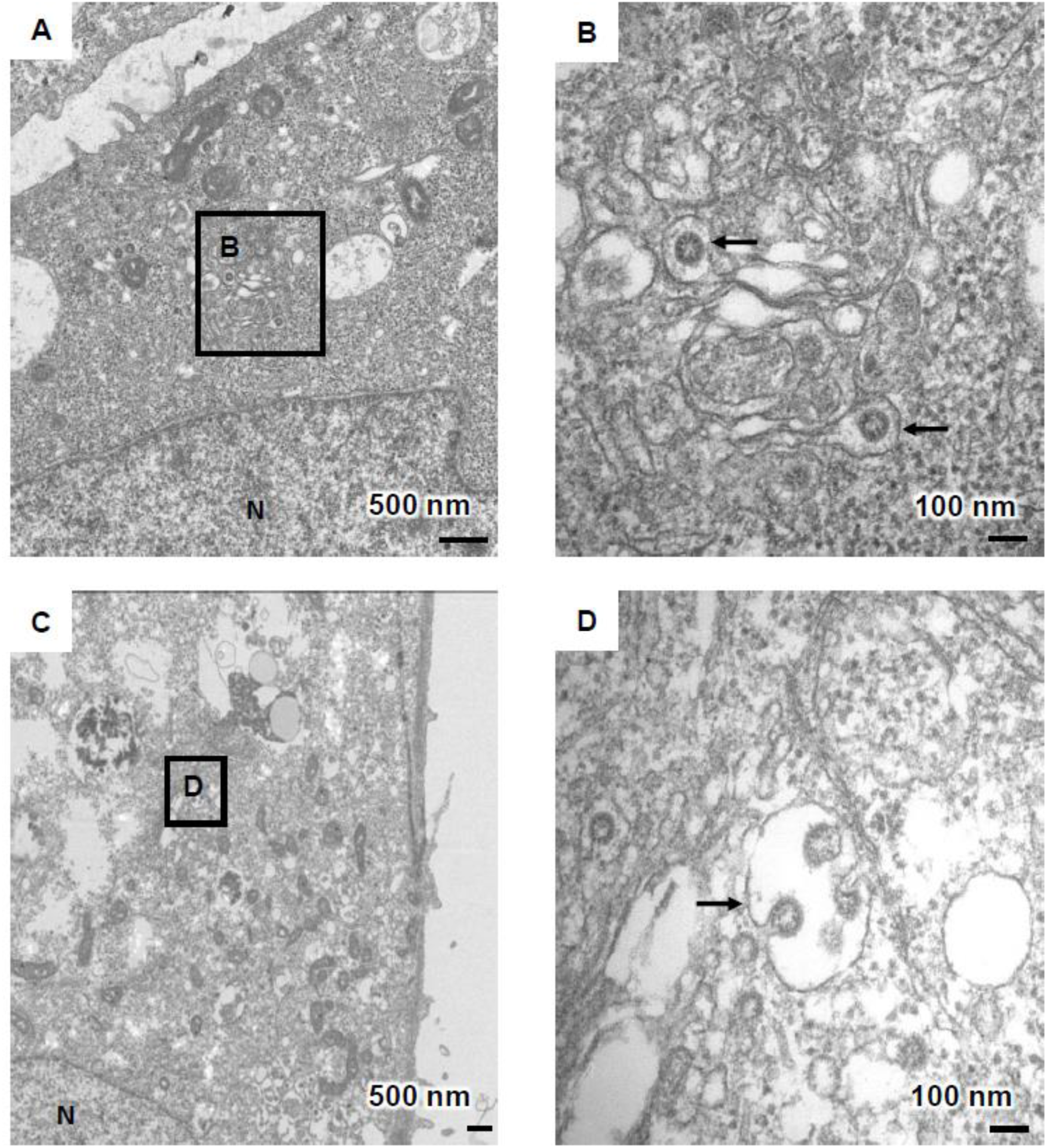
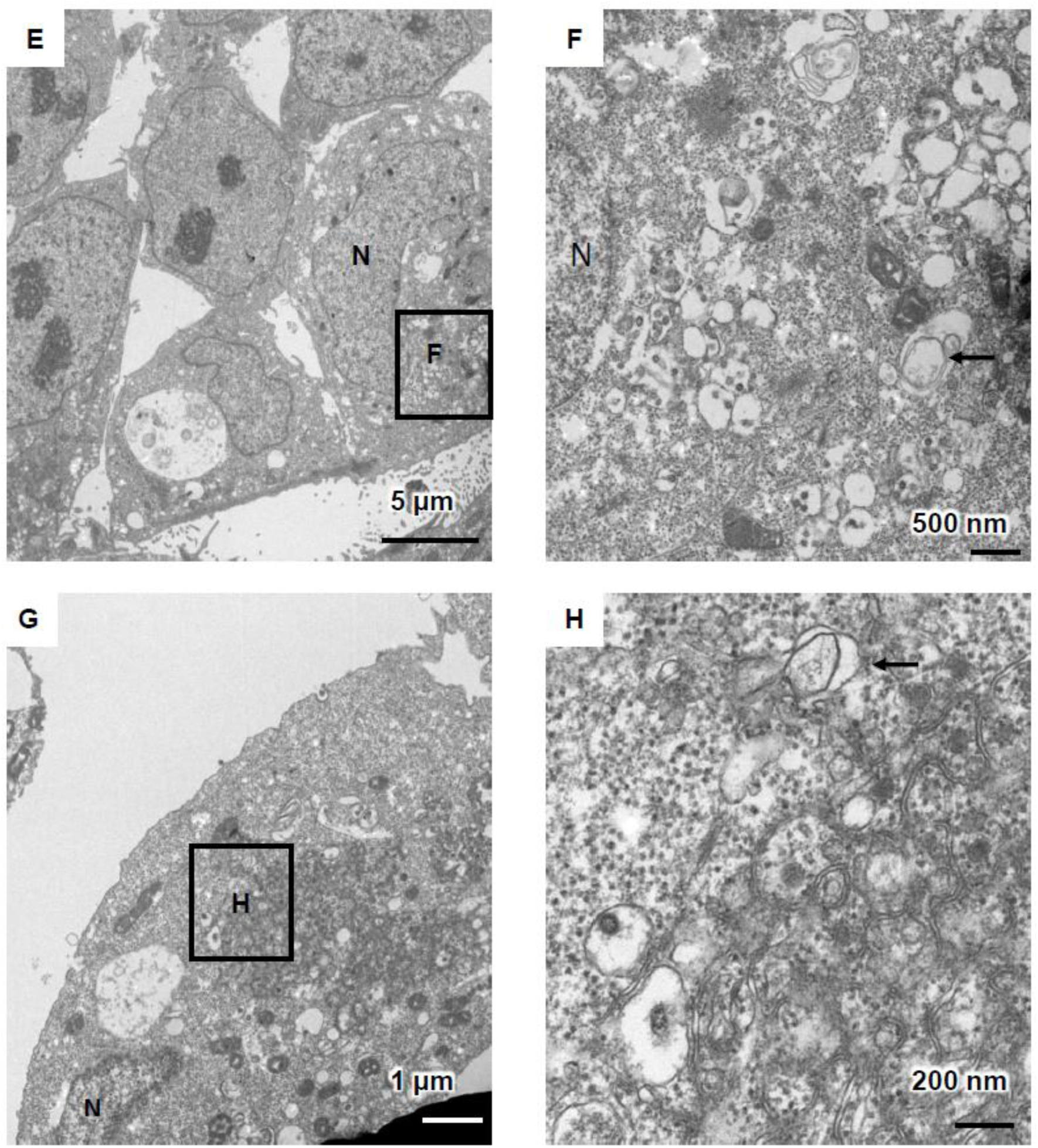
TEM images of infected ACE2-iPS cells. (**A-D**) Endoplasmic reticulum-Golgi intermediate compartment (ERGIC) containing SARS-CoV-2 particles (black arrows) was observed in infected ACE2-iPS cells. (**E-H**) Double membrane vesicles (black arrows) were observed in infected ACE2-iPS cells.

**Figure S4.**
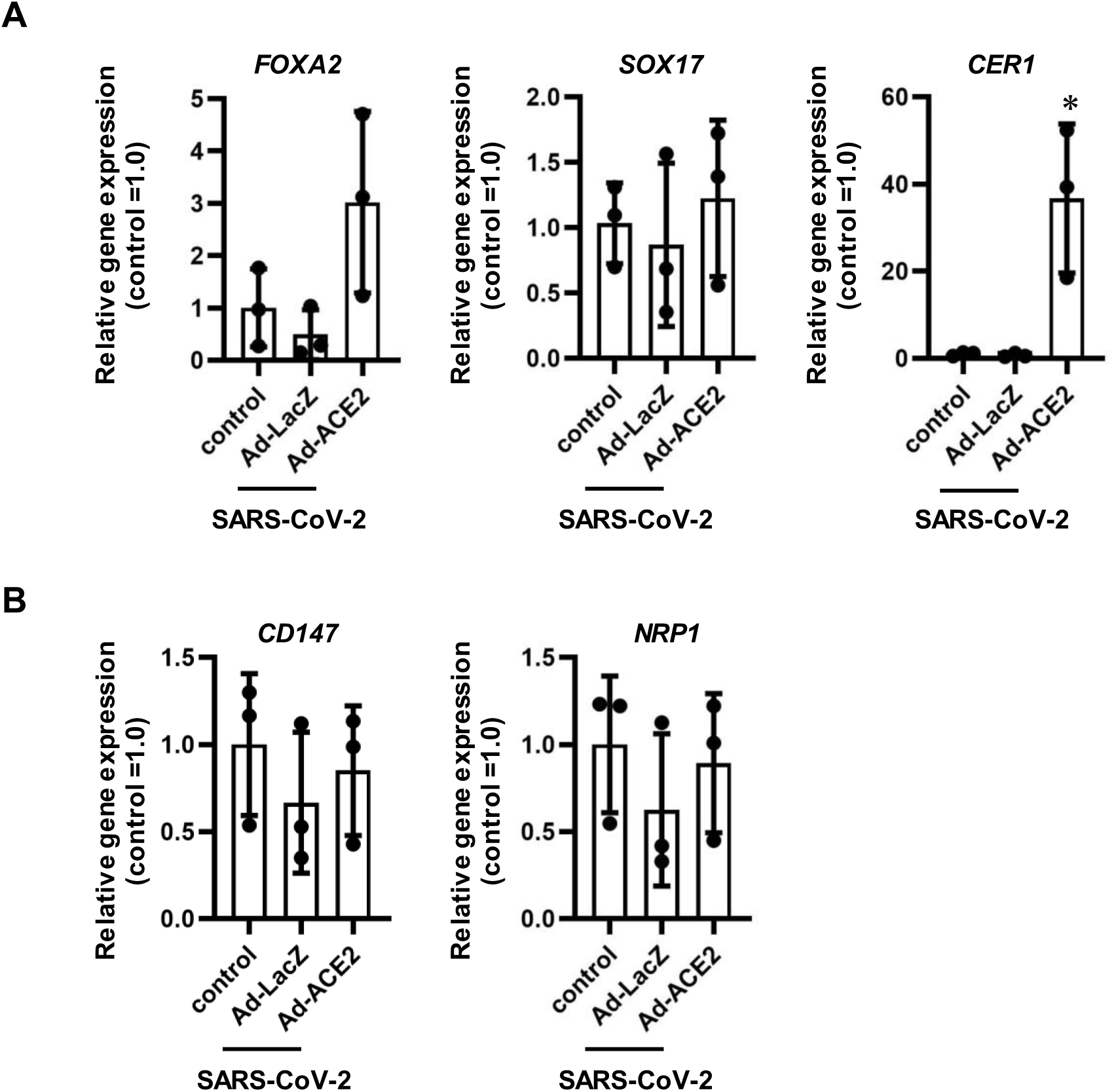
Gene expression profiles of differentiation markers and viral receptors in infected ACE2-iPS cells. Undifferentiated human iPS cells (1383D6) were transduced with 600 VP/cell of LacZ- or ACE2-expressing Ad vectors (Ad-LacZ and Ad-ACE2, respectively) for 2 hr and then cultured with AK02 medium for 2 days. Control human iPS cells were not transduced with Ad vectors. The LacZ- and ACE2-expressing human iPS cells were then infected with SARS-CoV-2 (5×10^4^ TCID50/well) for 2 hr and cultured with AK02 medium for 3 days. (**A, B**) The gene expression levels of endoderm markers (*FOXA2*, *SOX17*, and *CER1*) (**A**) and viral receptors (*CD147* and *NRP1*) (**B**) were examined by qPCR. Data are shown as means ± SD (*n*=3). One-way ANOVA followed by Tukey’s post hoc test (\**p*<0.05, compared with Ad-LacZ).

**Figure S5.**
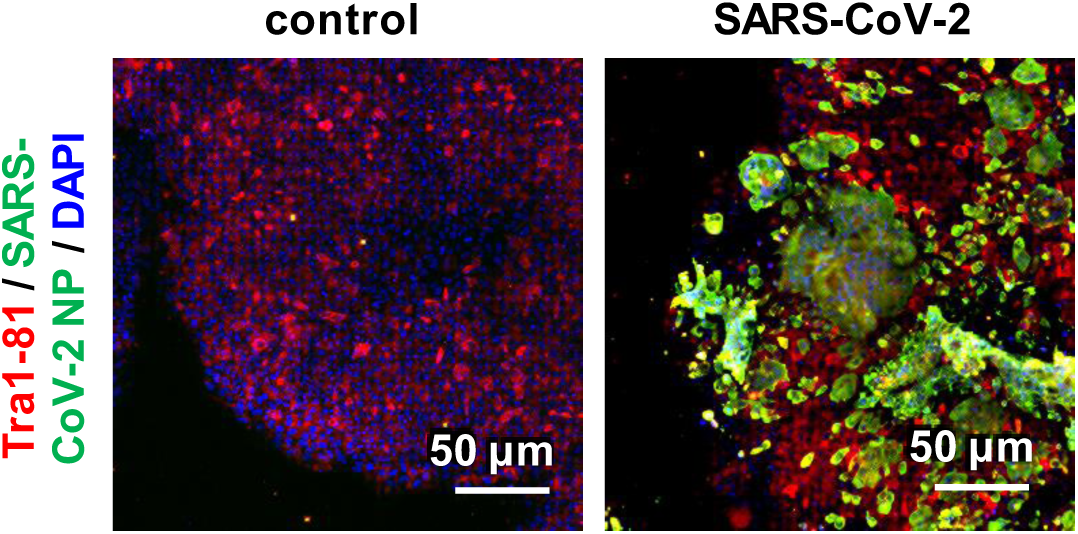
Immunofluorescence analysis of infected ACE2-iPS cells. Immunofluorescence analysis of SARS-CoV-2 NP (green) and OCT3/4 (red) in uninfected and infected ACE2-iPS cells. Nuclei were counterstained with DAPI (blue).

**Figure S6.**
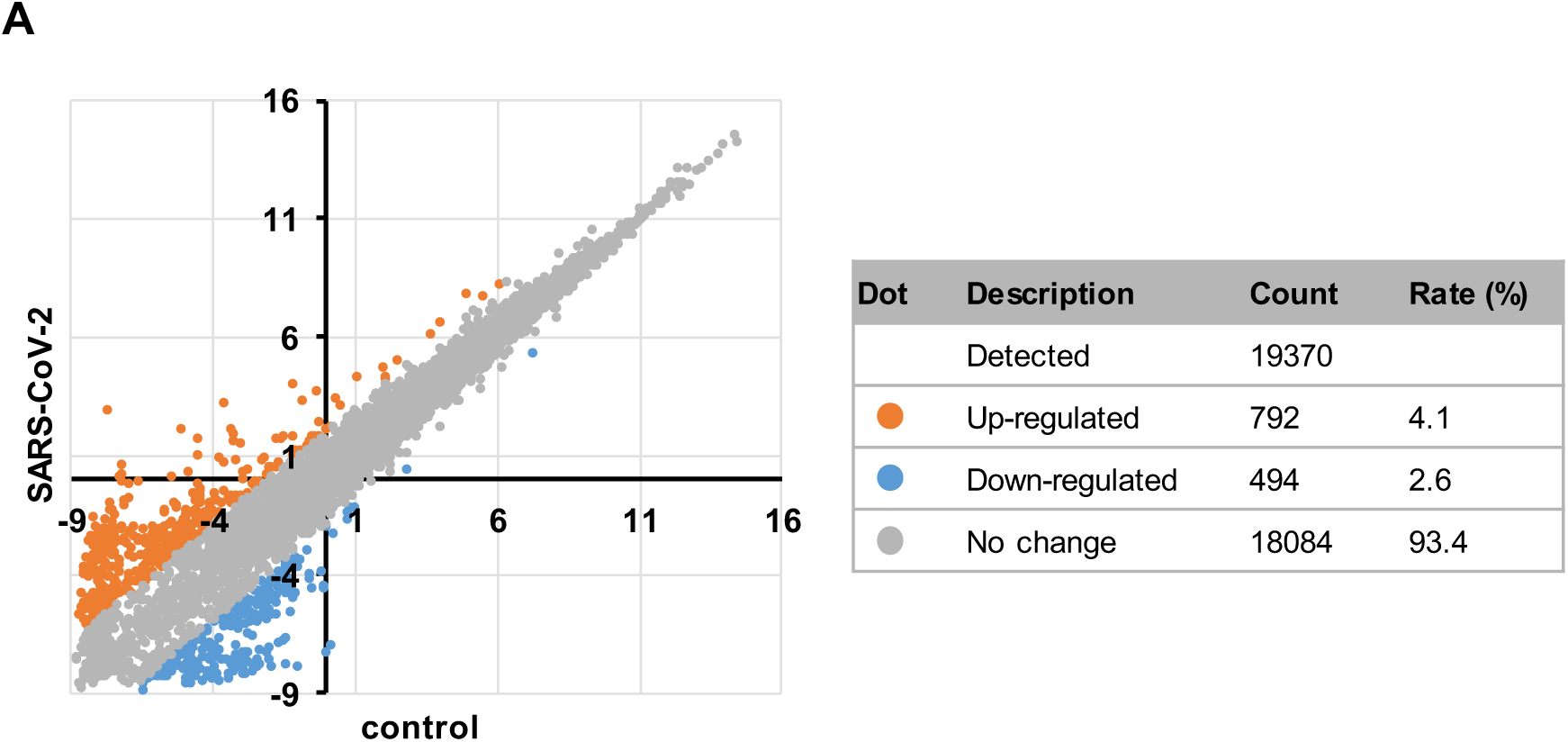
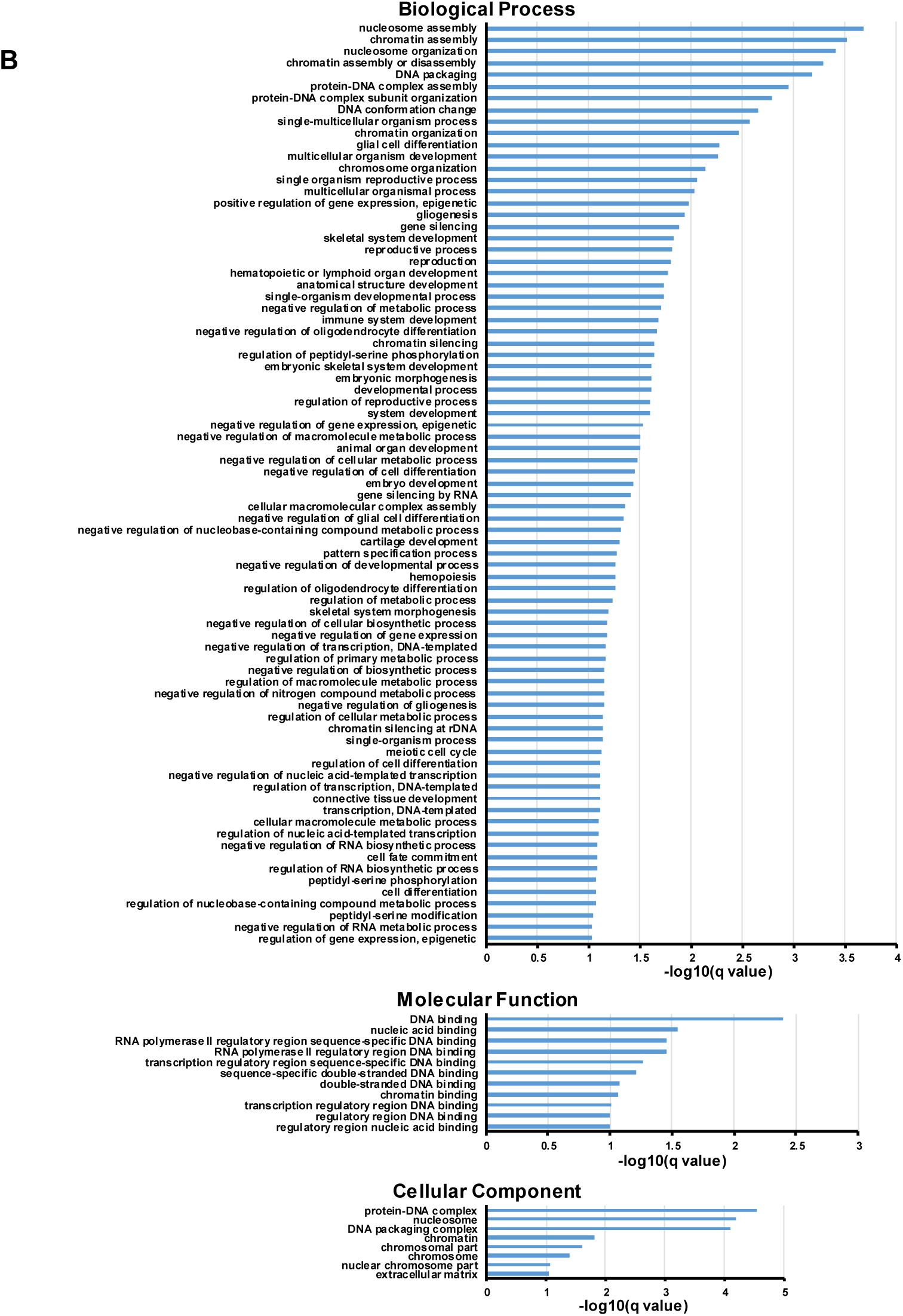
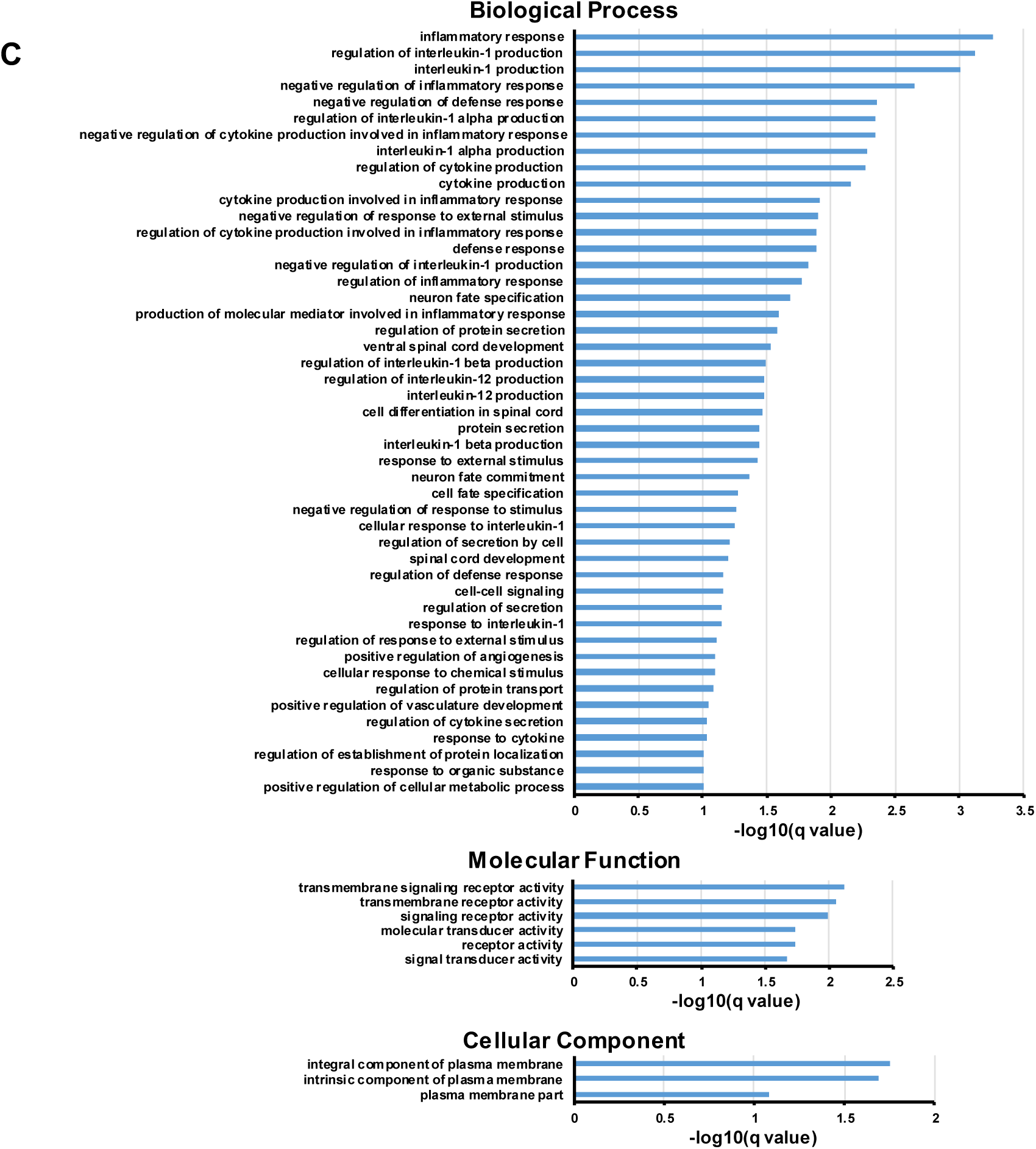
global gene expression analysis in infected ACE2-iPS cells. RNA-seq analysis was performed in uninfected and infected ACE2-iPS cells. (**A**) A scatter plot of uninfected ACE2-iPS cells (control) vs. infected ACE2-iPS cells (SARS-CoV-2). The count and rate of up-regulated and down-regulated genes are summarized in the table. (**B, C**) GO analysis was performed for gene sets whose gene expression levels were decreased (**B**) or increased (**C**) more than three-fold.

**Figure S7.**
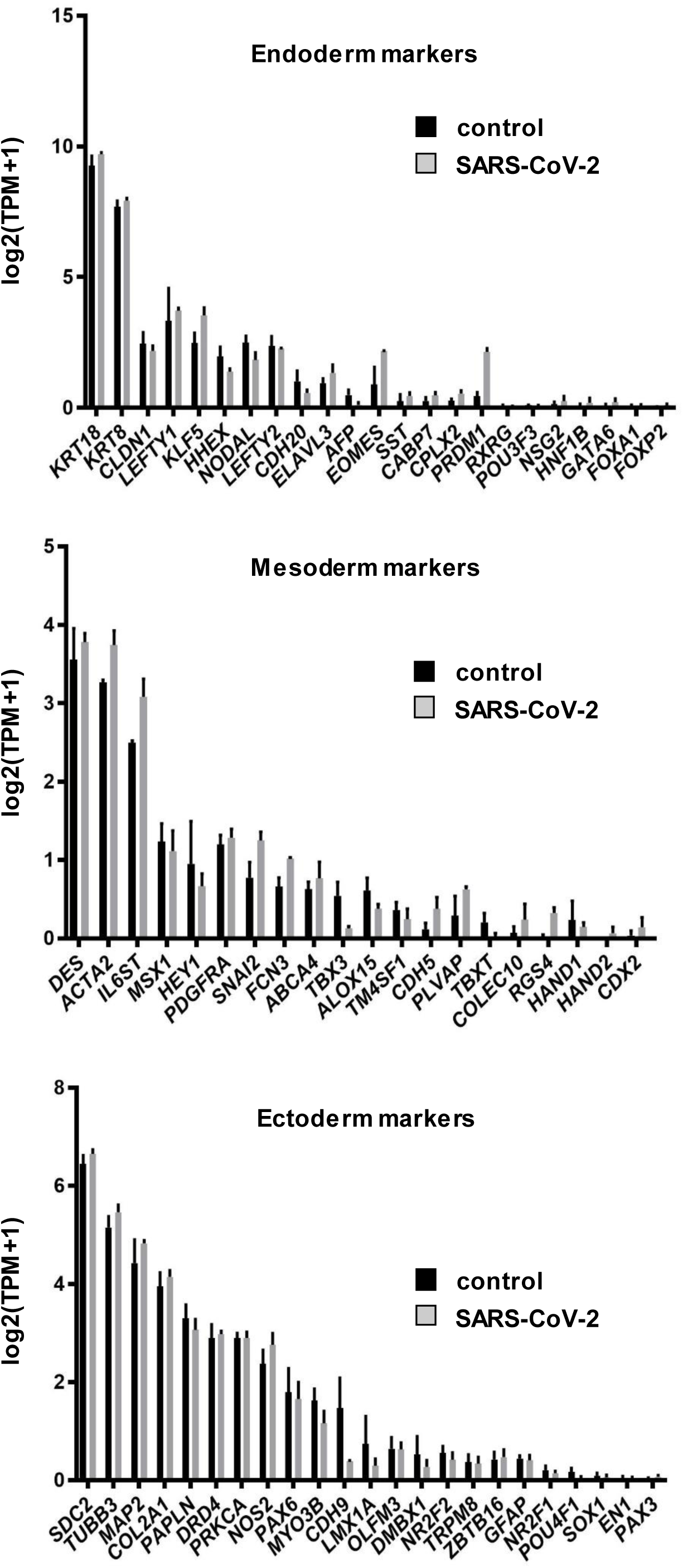
Gene expression profiles of endoderm, mesoderm, and ectoderm markers in infected ACE2-iPS cells. RNA-seq analysis was performed in uninfected and infected ACE2-iPS cells. Bar plots of ectoderm, mesoderm, and endoderm markers in uninfected ACE2-iPS cells (control) and infected ACE2-iPS cells (SARS-CoV-2) are shown.

**Figure S8.**
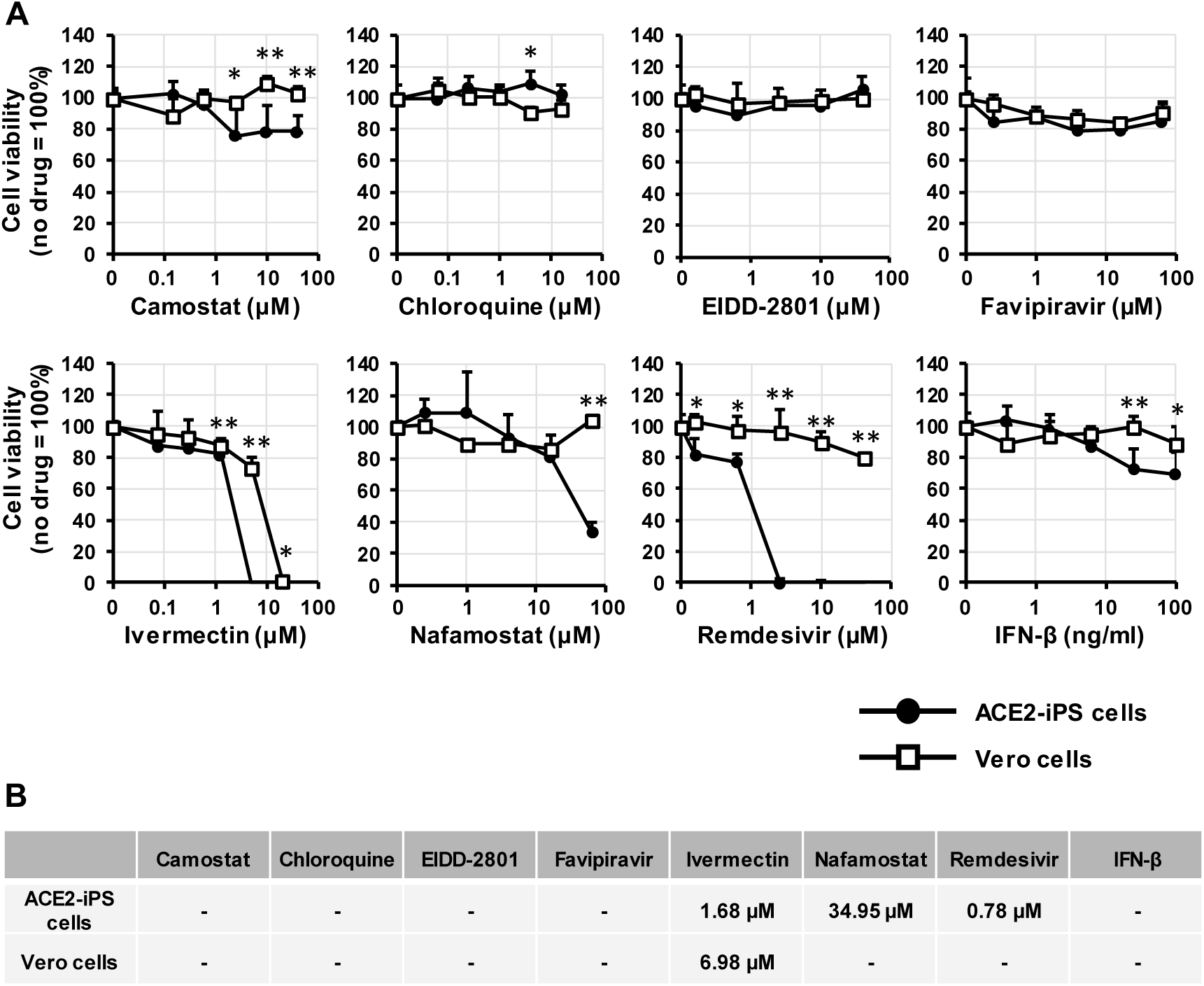
Cytotoxicity of anti-COVID-19 drugs in ACE2-iPS cells. (**A**) Vero cells and ACE2-iPS cells were cultured with medium containing drugs for 4 days. Cell viability was measured by the WST-8 assay. (**B**) The CC50 values (μM) of anti-COVID-19 drugs in Vero cells and ACE2-iPS cells were calculated using GraphPad Prism8 and summarized. Data are shown as means ± SD (*n*=3). Unpaired two-tailed Student’s *t*-test (\**p*<0.05, \*\**p*<0.01; ACE2-iPS vs Vero cells).

**Figure S9.**
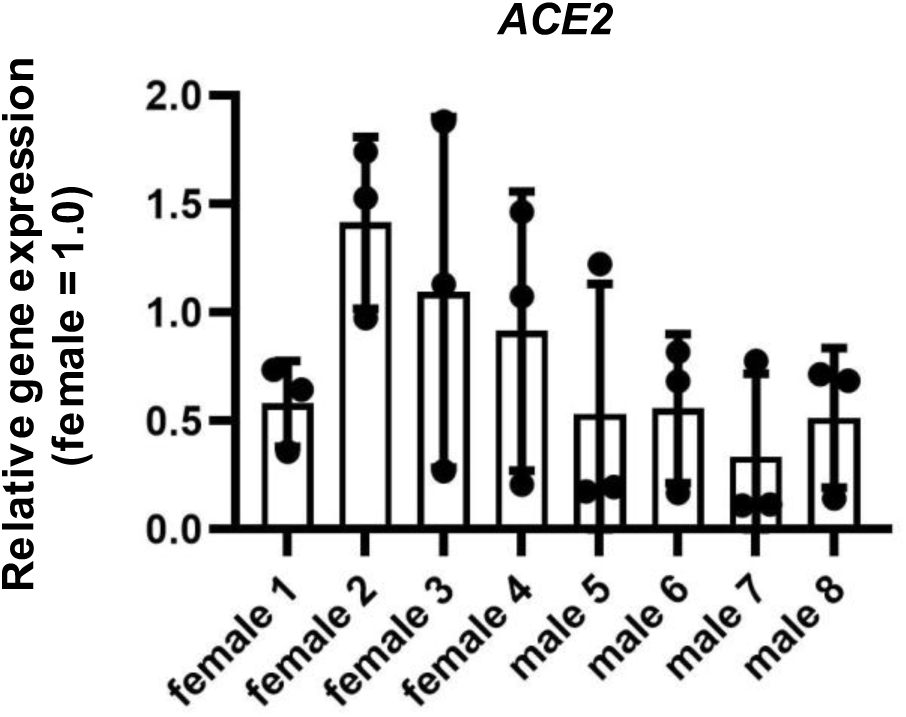
The *ACE2* expression levels in the female and male ACE2-iPS/ES cell lines. Four female ES/iPS cell lines and four male ES/iPS cell lines were transduced with 600 VP/cell of ACE2-expressing Ad vectors (Ad-ACE2) for 2 hr and then cultured with AK02 medium for 2 days. The *ACE2* expression levels were measured by qPCR for each cell line. Unpaired two-tailed Student’s *t*-test (\*\**p*<0.01). Data are shown as means ± SD (*n*=3). Female1:H9, Female2: KhES1, Female3:KhES2, Female4: 201B7, Male1:H1, Male2:KhES3, Male3:Tic, Male4:1383D6.

## Materials and Methods

**All unique/stable reagents generated in this study are available from the corresponding authors with a completed Materials Transfer Agreement.**

### Human ES/iPS cells

The human ES/iPS cell lines 1383D6 (Nakagawa et al., 2014) (provided by Dr. Masato Nakawaga, Kyoto University), 201B7 (Takahashi et al., 2007), Tic (JCRB1331, JCRB Cell Bank), H1 (WA01), H9 (WA09) (WiCell Research Institute), KhES1, KhES2, and KhES3 (Kyoto University) were maintained on 0.5 μg/cm^2^ recombinant human laminin 511 E8 fragments (iMatrix-511, Nippi) with StemFit AK02N medium (Ajinomoto) containing 10 μM Y-27632 (from day 0 to day 1, FUJIFILM Wako Pure Chemical). To passage human ES/iPS cells, near-confluent human ES/iPS cell colonies were treated with TrypLE Select Enzyme (Thermo Fisher Scientific) for 5 min at 37°C. After the centrifugation, the cells were seeded at an appropriate cell density (1.3×10^4^ cells/9 cm^2^) onto iMatrix-511 and subcultured every 6 days. Human ES cells were used following the Guidelines for Derivation and Utilization of Human Embryonic Stem Cells of the Ministry of Education, Culture, Sports, Science and Technology of Japan, and the study was approved by an independent ethics committee. Except for **figures 6 and S9**, a human iPS cell line, 1383D6, was used for all experiments.

### SARS-CoV-2 preparation

The SARS-CoV-2 strains used in this study (SARS-CoV-2/Hu/DP/Kng/19-027) were provided from the Kanagawa Prefectural Institute of Public Health. SARS-CoV-2 was isolated from a COVID-19 patient (GenBank: LC528233.1). The isolation and analysis of the virus will be described elsewhere (manuscript in preparation). The virus was plaque-purified and propagated in Vero cells and stored at −80°C. All experiments including virus infections were done in a biosafety level 3 facility at Kyoto University strictly following regulations.

### Adenovirus vectors

Ad vectors were constructed using Adeno-X^TM^ Adenoviral System 3 (Takara Bio). The ACE2 and TMPRSS2 genes were amplified by PCR using cDNA generated from Pulmonary Alveolar Epithelial Cell Total RNA (ScienCell Research Laboratories) as a template. The ACE2 and TMPRSS2 genes were inserted into Adeno-X^TM^ Adenoviral System 3, resulting in pAdX-ACE2 and pAdX-TMPRSS2, respectively. The ACE2- and TMPRSS2-expressing Ad vectors (Ad-ACE2 and Ad-TMPRSS2, respectively) were propagated in HEK293 cells (JCRB9068, JCRB Cell Bank). LacZ-expressing Ad vectors were purchased from Vector Biolabs. The vector particle (VP) titer was determined by using a spectrophotometric method (Maizel Jr et al., 1968).

### SARS-CoV-2 infection and drug treatment

ACE2-iPS cells and Vero cells (JCRB0111, JCRB Cell Bank) cultured in a 96-well plate (2.0×10^4^ cells/well) were infected with 2.0×10^3^ TCID50/well of SARS-CoV-2 for 2 hr and then replaced with medium containing drugs. In the infection and drug treatment experiments, the medium containing drugs was replaced with fresh medium every day. At day 2 (Vero cells) or day 4 (ACE2-iPS cells) after the infection, the viral RNA copy number in the cell culture supernatant was measured by qPCR. Drugs used in the infection experiments are summarized in **Table S1**.

**Table S1.** Drugs used in the infection experiments.

| Drug | Catalogue | Company |
| --- | --- | --- |
| Camostat mesylate | SML0057 | Sigma-Aldrich |
| Chloroquine diphosphate | No.4109 | Tocris |
| EIDD-2801 | HY-135853 | MedChemExpress |
| Favipiravir | S7975 | Selleck Chemicals |
| Ivermectin | I8618 | LKT Labs |
| Nafamostat mesylate | N0289-10MG | Sigma-Aldrich |
| Recombinant human IFN-β | 300-02BC | PeproTech |
| Remdesivir | A17170 | Clinisciences |

### Viral titration of SARS-CoV-2

Viral titers were measured by median tissue culture infectious dose (TCID50) assays at a biosafety level 3 laboratory (Kyoto University). TMPRSS2/Vero cells (JCRB1818, JCRB Cell Bank) (Matsuyama et al., 2020) were cultured with Minimum Essential Media (MEM, Sigma-Aldrich) supplemented with 5% fetal bovine serum (FBS), and 1% penicillin/streptomycin and seeded into 96-well cell culture plates (Thermo Fisher Scientific). The samples were serially diluted 10-fold from 10^−1^ to 10^−8^ in the cell culture medium. Dilutions were placed onto the TMPRSS2/Vero cells in triplicate and incubated at 37°C for 96 hr. Cytopathic effects were evaluated under a microscope. TCID50/mL was calculated using the Reed-Muench method.

### Quantification of viral RNA copy number

The cell culture supernatant was mixed with an equal volume of 2×RNA lysis buffer (distilled water containing 0.4 U/uL SUPERase I^TM^ RNase Inhibitor (Thermo Fisher Scientific), 2% Triton X-100, 50 mM KCl, 100 mM Tris-HCl (pH 7.4), and 40% glycerol) and incubated at room temperature for 10 min. The mixture was diluted 10 times with distilled water. Viral RNA was quantified using a One Step TB Green PrimeScript PLUS RT-PCR Kit (Perfect Real Time) (Takara Bio) on a StepOnePlus real-time PCR system (Thermo Fisher Scientific). The primers used in this experiment are as follows: (forward) AGCCTCTTCTCGTTCCTCATCAC and (reverse) CCGCCATTGCCAGCCATTC. Standard curves were prepared using SARS-CoV-2 RNA (10^5^ copies/μL) purchased from Nihon Gene Research Laboratories.

### Quantitative PCR

Total RNA was isolated from human iPS cells using ISOGENE (NIPPON GENE). cDNA was synthesized using 500 ng of total RNA with a Superscript VILO cDNA Synthesis Kit (Thermo Fisher Scientific). Real-time RT-PCR was performed with SYBR Green PCR Master Mix (Thermo Fisher Scientific) using a StepOnePlus real-time PCR system (Thermo Fisher Scientific). The relative quantitation of target mRNA levels was performed by using the 2^-ΔΔCT^ method. The values were normalized by those of the housekeeping gene, *glyceraldehyde 3-phosphate dehydrogenase* (*GAPDH*). The PCR primer sequences are shown in **Table S2**.

**Table S2.**
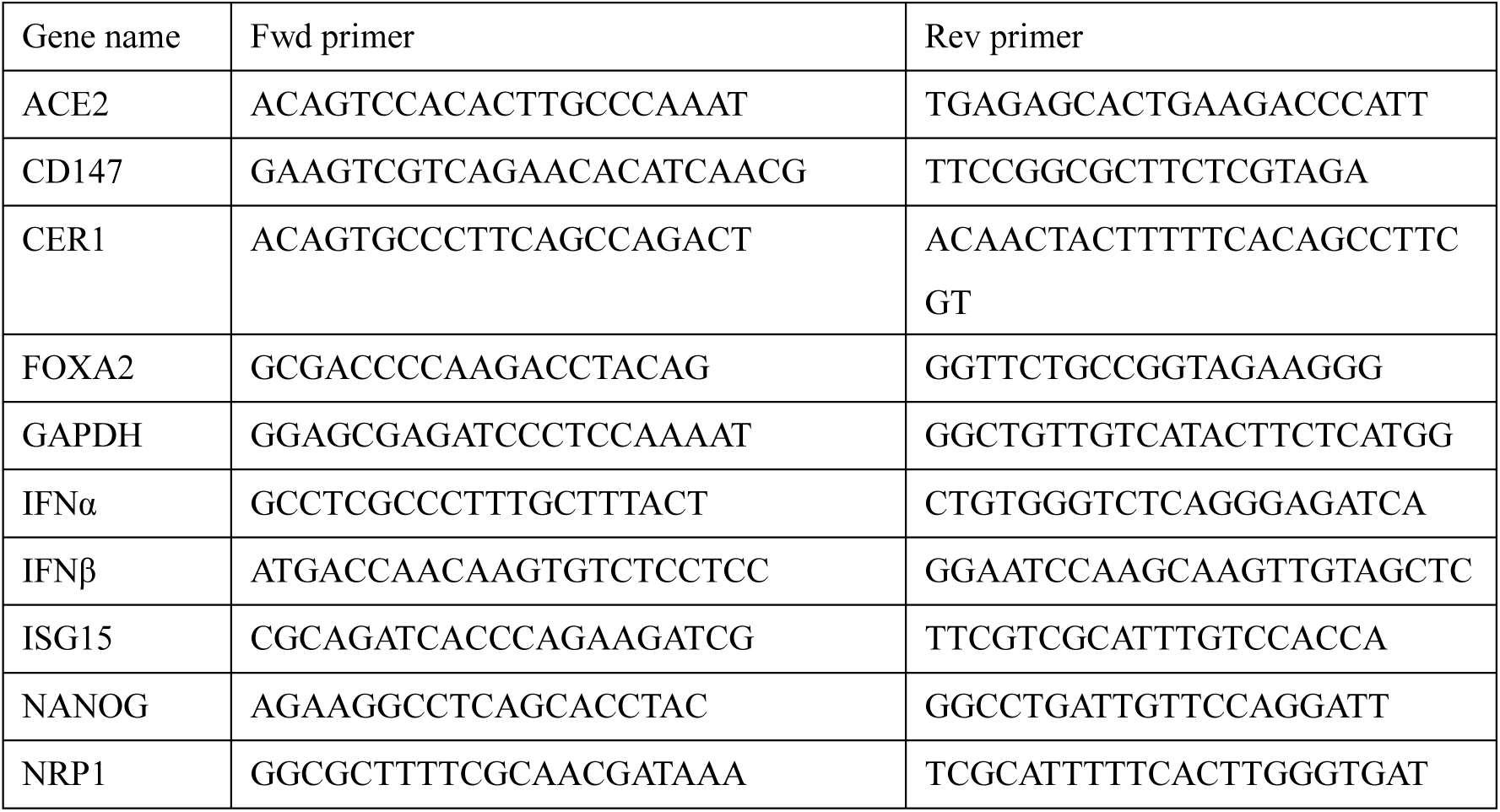
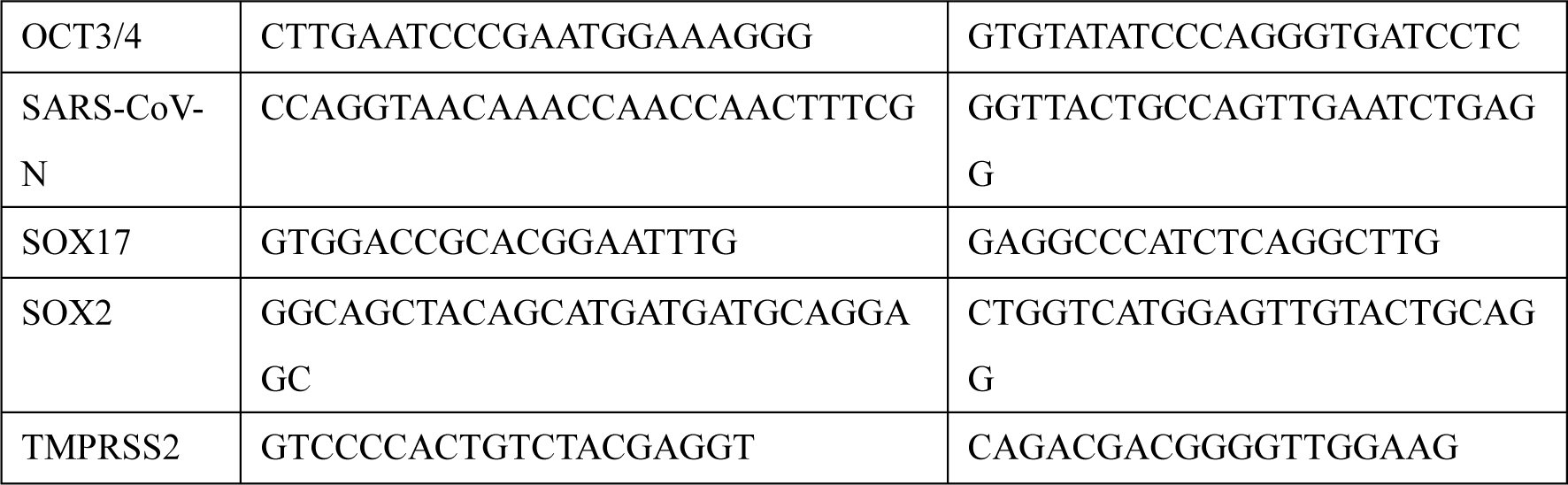
Primers used in this study.

### Ultrathin section transmission electron microscopy (TEM)

Uninfected and infected ACE2-iPS cells were fixed in phosphate buffer with 2% glutaraldehyde and subsequently post-fixed in 2% osmium tetra-oxide for 2 hr at 4°C. After fixation, the cells were dehydrated in a graded series of ethanol and embedded in epoxy resin. Ultrathin sections were cut, stained with uranyl acetate and lead staining solution, and examined using an electron microscope (HITACHI H-7600) at 100 kV.

### Immunofluorescence staining

For the immunofluorescence staining of human iPS cells, the cells were fixed with 4% paraformaldehyde in PBS at 4°C. After blocking the cells with PBS containing 2% bovine serum albumin and 0.2% Triton X-100 at room temperature for 45 min, the cells were incubated with a primary antibody at 4°C overnight and then with a secondary antibody at room temperature for 1 hr. All antibodies used in this report are described in **Table S3**.

**Table S3.** Antibodies used in this study.

| Antigen | Catalogue | Host | Company |
| --- | --- | --- | --- |
| Donkey anti-Mouse IgG (H+L)<br>Highly Cross-Adsorbed Secondary Antibody,<br>Alexa Fluor Plus 594 | A32744 | Donkey | Thermo Fisher Scientific |
| Goat anti-Rabbit IgG (H+L)<br>Highly Cross-Adsorbed Secondary Antibody,<br>Alexa Fluor Plus 488 | A32731 | Goat | Thermo Fisher Scientific |
| OCT3/4 | sc-5279 | Mouse | Santa Cruz Biotechnology |
| SARS-CoV-2 NP | A2061-50 | Rabbit | BIO Vision |
| SOX2 | sc-365823 | Mouse | Santa Cruz Biotechnology |
| Tra1-81 | MAB4381 | Mouse | Sigma-Aldrich |

### RNA-seq

Total RNA was prepared using the RNeasy Mini Kit (Qiagen). RNA integrity was assessed with a 2100 Bioanalyzer (Agilent Technologies). The library preparation was performed using a TruSeq stranded mRNA sample prep kit (Illumina) according to the manufacturer’s instructions. Sequencing was performed on an Illumina NextSeq500. The fastq files were generated using bcl2fastq-2.20. Adapter sequences and low-quality bases were trimmed from the raw reads by Cutadapt ver 1.14 (Martin, 2011). The trimmed reads were mapped to the human reference genome sequences (hg38) using STAR ver 2.5.3a (Dobin et al., 2013) with the GENCODE (release 36, GRCh38.p13) (Frankish et al., 2019) gtf file. The raw counts for protein-coding genes were calculated using htseq-count ver 0.12.4 (Anders et al., 2015) with the GENCODE gtf file. Gene expression levels were determined as transcripts per million (TPM) with DEseq2 (Love et al., 2014). Raw data concerning this study were submitted under Gene Expression Omnibus (GEO) accession number GSE166990.

### Statistical analyses

Statistical significance was evaluated by unpaired two-tailed Student’s *t*-test or one-way analysis of variance (ANOVA) followed by Tukey’s post hoc tests. Statistical analyses were performed using GraphPad Prism8 and 9. Data are representative of three independent experiments. Details are described in the figure legends.

## References

Anastassopoulou, C., Gkizarioti, Z., Patrinos, G.P., and Tsakris, A. (2020). Human genetic factors associated with susceptibility to SARS-CoV-2 infection and COVID-19 disease severity. Human genomics 14, 1–8.

Anders, S., Pyl, P.T., and Huber, W. (2015). HTSeq—a Python framework to work with high-throughput sequencing data. Bioinformatics 31, 166–169.

Dobin, A., Davis, C.A., Schlesinger, F., Drenkow, J., Zaleski, C., Jha, S., Batut, P., Chaisson, M., and Gingeras, T.R. (2013). STAR: ultrafast universal RNA-seq aligner. Bioinformatics 29, 15–21.

Frankish, A., Diekhans, M., Ferreira, A.-M., Johnson, R., Jungreis, I., Loveland, J., Mudge, J.M., Sisu, C., Wright, J., and Armstrong, J. (2019). GENCODE reference annotation for the human and mouse genomes. Nucleic acids research 47, D766–D773.

Group, S.C.-G. (2020). Genomewide association study of severe Covid-19 with respiratory failure. New England Journal of Medicine 383, 1522–1534.

Han, Y., Duan, X., Yang, L., Nilsson-Payant, B.E., Wang, P., Duan, F., Tang, X., Yaron, T.M., Zhang, T., and Uhl, S. (2021). Identification of SARS-CoV-2 inhibitors using lung and colonic organoids. Nature 589, 270–275.

Hoffmann, M., Kleine-Weber, H., Schroeder, S., Krüger, N., Herrler, T., Erichsen, S., Schiergens, T.S., Herrler, G., Wu, N.-H., and Nitsche, A. (2020). SARS-CoV-2 cell entry depends on ACE2 and TMPRSS2 and is blocked by a clinically proven protease inhibitor. cell 181, 271–280.e278.

Kajiwara, M., Aoi, T., Okita, K., Takahashi, R., Inoue, H., Takayama, N., Endo, H., Eto, K., Toguchida, J., and Uemoto, S. (2012). Donor-dependent variations in hepatic differentiation from human-induced pluripotent stem cells. Proceedings of the National Academy of Sciences 109, 12538–12543.

Kim, S.-I., Matsumoto, T., Kagawa, H., Nakamura, M., Hirohata, R., Ueno, A., Ohishi, M., Sakuma, T., Soga, T., and Yamamoto, T. (2018). Microhomology-assisted scarless genome editing in human iPSCs. Nature communications 9, 1–14.

Klein, S., Cortese, M., Winter, S.L., Wachsmuth-Melm, M., Neufeldt, C.J., Cerikan, B., Stanifer, M.L., Boulant, S., Bartenschlager, R., and Chlanda, P. (2020). SARS-CoV-2 structure and replication characterized by in situ cryo-electron tomography. Nature communications 11, 1–10.

Love, M.I., Huber, W., and Anders, S. (2014). Moderated estimation of fold change and dispersion for RNA-seq data with DESeq2. Genome biology 15, 1–21.

Maier, H.J., Hawes, P.C., Cottam, E.M., Mantell, J., Verkade, P., Monaghan, P., Wileman, T., and Britton, P. (2013). Infectious bronchitis virus generates spherules from zippered endoplasmic reticulum membranes. MBio 4.

Maizel Jr, J.V., White, D.O., and Scharff, M.D. (1968). The polypeptides of adenovirus: I. Evidence for multiple protein components in the virion and a comparison of types 2, 7A, and 12. Virology 36, 115–125.

Martin, M. (2011). Cutadapt removes adapter sequences from high-throughput sequencing reads. EMBnet journal 17, 10–12.

Matsuyama, S., Nao, N., Shirato, K., Kawase, M., Saito, S., Takayama, I., Nagata, N., Sekizuka, T., Katoh, H., and Kato, F. (2020). Enhanced isolation of SARS-CoV-2 by TMPRSS2-expressing cells. Proceedings of the National Academy of Sciences 117, 7001–7003.

Nakagawa, M., Taniguchi, Y., Senda, S., Takizawa, N., Ichisaka, T., Asano, K., Morizane, A., Doi, D., Takahashi, J., and Nishizawa, M. (2014). A novel efficient feeder-free culture system for the derivation of human induced pluripotent stem cells. Scientific reports 4, 1–7.

Pairo-Castineira, E., Clohisey, S., Klaric, L., Bretherick, A.D., Rawlik, K., Pasko, D., Walker, S., Parkinson, N., Fourman, M.H., and Russell, C.D. (2020). Genetic mechanisms of critical illness in Covid-19. Nature, 1-1.

Park, I.-H., Arora, N., Huo, H., Maherali, N., Ahfeldt, T., Shimamura, A., Lensch, M.W., Cowan, C., Hochedlinger, K., and Daley, G.Q. (2008). Disease-specific induced pluripotent stem cells. cell 134, 877–886.

Takahashi, K., Tanabe, K., Ohnuki, M., Narita, M., Ichisaka, T., Tomoda, K., and Yamanaka, S. (2007). Induction of pluripotent stem cells from adult human fibroblasts by defined factors. cell 131, 861–872.

Takayama, K. (2020). In vitro and animal models for SARS-CoV-2 research. Trends in pharmacological sciences 41, 513–517.

Wambier, C.G., Goren, A., Vaño-Galván, S., Ramos, P.M., Ossimetha, A., Nau, G., Herrera, S., and McCoy, J. (2020). Androgen sensitivity gateway to COVID-19 disease severity. Drug development research 81, 771–776.

Wang, K., Chen, W., Zhang, Z., Deng, Y., Lian, J.-Q., Du, P., Wei, D., Zhang, Y., Sun, X.-X., and Gong, L. (2020). CD147-spike protein is a novel route for SARS-CoV-2 infection to host cells. Signal transduction and targeted therapy 5, 1–10.

Weiss, P., and Murdoch, D.R. (2020). Clinical course and mortality risk of severe COVID-19. The Lancet 395, 1014–1015.

Westmeier, J., Paniskaki, K., Karaköse, Z., Werner, T., Sutter, K., Dolff, S., Overbeck, M., Limmer, A., Liu, J., and Zheng, X. (2020). Impaired cytotoxic CD8+ T cell response in elderly COVID-19 patients. MBio 11.

Zeberg, H., and Pääbo, S. (2020). The major genetic risk factor for severe COVID-19 is inherited from Neanderthals. Nature 587, 610–612.

Ziegler, C.G., Allon, S.J., Nyquist, S.K., Mbano, I.M., Miao, V.N., Tzouanas, C.N., Cao, Y., Yousif, A.S., Bals, J., and Hauser, B.M. (2020). SARS-CoV-2 receptor ACE2 is an interferon-stimulated gene in human airway epithelial cells and is detected in specific cell subsets across tissues. Cell 181, 1016–1035.e1019.

